# CD34 serves as an intrinsic innate immune guardrail protecting stem cells from replicating retroviruses

**DOI:** 10.1101/2025.03.15.643450

**Authors:** Sijia He, Amrita Haikerwal, Sameer Tiwari, Deemah Dabbagh, Mohammed Z. Alam, Janice L. Yoon, Brian Hetrick, Yang Han, Liang Shan, Christopher Lockhart, Yuntao Wu

**Author notes:** These authors contributed equally to this work (Sijia He, Amrita Haikerwal, Sameer Tiwari, Deemah Dabbagh). Vibrant Biomedicines, Inc., Cambridge, MA 02142, USA. Virongy Biosciences, Inc., Manassas, VA 20109, USA. **Correspondence:** (Y.W).

## Abstract

Stem cells are highly resistant to viral infection compared to their differentiated progeny, and this resistance is associated with stem cell-specific restriction factors and intrinsic interferon stimulated genes (ISGs). In HIV infection, proviral DNA has been detected in certain bone marrow hematopoietic stem cells, yet widespread stem cell infection *in vivo* is restricted. Intriguingly, exposing bone marrow stem cells to HIV *in vitro* led to viral replication selectively only in the CD34^-^ population, but not in the CD34^+^ cells. The mechanism dictating this CD34-based HIV restriction remained a mystery, especially since HIV has a capacity to antagonize restriction factors and ISGs. CD34 is a common marker of hematopoietic stem and progenitor cells. Here, we report the intrinsic antiviral properties of CD34. Expression of CD34 in HIV-1 producer cells results in the loss of progeny virion infectivity. Conversely, removal of CD34 using CRISPR/Cas9 knockout or stem cell differentiation cytokines promotes HIV-1 replication in stem cells. These results suggest that in addition to restriction factors and intrinsic ISGs, CD34 serves as a host innate protection preventing retrovirus replication in stem cells. Mechanistically, CD34 does not block viral entry, integration, and release. Instead, it becomes incorporated onto progeny virions, which inactivates virus infectivity. These findings offer new insights into innate immunity in stem cells, and highlight intriguing retrovirus-host interactions in evolution.

## Introduction

Uncontrolled retroviral proliferation and genome integration in stem and progenitor cells can lead to the interference of stem cell maturation and interruption of normal hematopoiesis, a detrimental condition that can result in multilineage hematopoietic failure and malignancy ^1^. Nevertheless, it has long been recognized that stem cells are highly resistant to viral infection compared to their differentiated progeny ^2-6^, and this resistance is associated with specific expression of restriction factors and intrinsic interferon stimulated genes (ISGs) ^3,7^. In human immunodeficiency virus (HIV) infection, the infection of undifferentiated stem and progenitor cells is also limited, and frequently results in low-level viral production or latency that requires differentiation factors for active viral replication ^1,8-10^. For example, it has been shown that HIV rarely infects bone marrow hematopoietic stem and progenitor cells *in vivo* ^11-13^, although some studies have detected HIV-DNA in the bone marrow progenitor cells from HIV patients ^10,14^. *In vitro*, direct HIV infection of bone marrow hematopoietic progenitor cells or immature CD34^+^CD38^-^ stem cells also did not result in detectable HIV replication ^5,15^, whereas infection of CD34^+^CD38^+^ bone marrow stem cells only led to low-level, limited HIV replication ^5^; low-level viral production was also observed after 40 to 60 days of HIV infection of bone marrow progenitor cells, when these cells were phenotypically differentiated into CD4^+^ monocytes ^16^. Intriguingly, exposure of HIV to bone marrow stem cells led to HIV replication selectively only in the CD34^-^ cell population, but not in the CD34^+^ cell population ^17^. The underlying mechanism dictating this CD34-based HIV selectivity remained mysterious, especially given that HIV has evolved accessory proteins such as Vif, Vpu, and Nef to antagonize restriction factors and ISGs ^18-28^.

CD34 is a transmembrane phosphoglycoprotein originally identified on hematopoietic stem cells ^29,30^ and on the progenitor cells of a variety of tissues ^31^. Expression of CD34 is generally associated with the phenotypes of hematopoietic cells and multiple nonhematopoietic progenitor cells, and thus serves as a common marker of diverse progenitor cells ^31^. Nevertheless, little is known about the exact function of CD34, although the molecule has been suggested to facilitate cell adhesion and migration ^32-36^ or, under some circumstances, block cell adhesion ^37,38^. CD34 is also a member of the sialomucin family of proteins, and was found to bind to L-selectin, P-selectin, and E-selectins ^39,40^. A shared ligand for these selectins, PSGL-1, has recently been identified as an anti-HIV restriction factor, inactivating virus infectivity through virion incorporation that inhibits virion attachment to target cells ^26-28^. In this study, we investigated whether CD34 has a similar antiviral activity in stem and progenitor cells, using HIV-1 infection as a model. Our results demonstrate that expression of CD34 in human CD4 T cells or undifferentiated CD34^+^ lymphoblastic progenitor Kasumi-3 cells inhibited HIV-1 replication, whereas CRISPR/Cas9 knockout of CD34 in the progenitor cell promoted HIV replication. We further demonstrated that direct infection of primary human CD34^+^ cord blood hematopoietic stem and progenitor cells with a VSV-G-pseudotyped HIV-1 led to the release of non-infectious virus, whereas treatment of the cells with the stem cell differentiation cytokines, GM-CSF and INF-α, downregulated CD34 and promoted the production of infectious HIV-1. Mechanistically, cell surface CD34 does not block viral entry and release. Instead, it is incorporated onto progeny virions during viral assembly, thereby inactivating virus infectivity through blocking particle attachment to target cells. These results suggest that ubiquitous expression of CD34 on stem and progenitor cells serves as a host innate immune defense limiting retroviral dissemination in these critical cell populations.

## Results

### Virion Incorporation of CD34 Inactivates HIV-1 Infectivity

CD34 is a sialomucin with the extracellular domain (NC) that is heavily glycosylated ^41^. We modelled the structure of the CD34 NC (residues 32-290) using Rosetta ^42-46^. The initial structure was built as a linear chain before core-1 or core-2 O-glycans were added to Thr/Ser residues, and core N-glycans were added to Asn residues, satisfying the N-X-S/T motif. Because O-glycans were added randomly, 100 such structures were built and subjected to relaxation. We compared these results to structures where no glycans were added, and found that with glycans, the end-to-end distance of CD34 is 226 ± 89 Å, and without glycans, the distance increases to 260 ± 110 Å. A representation of one glycosylated CD34 structure is presented in **Figure 1A**. Our structural analysis indicates that CD34 glycans, which are predominately localized to the CD34 N-terminus situated away from the cell membrane, have a condensing effect that may serve to increase the glycan density of CD34 on the cell surface.

**Figure 1.**
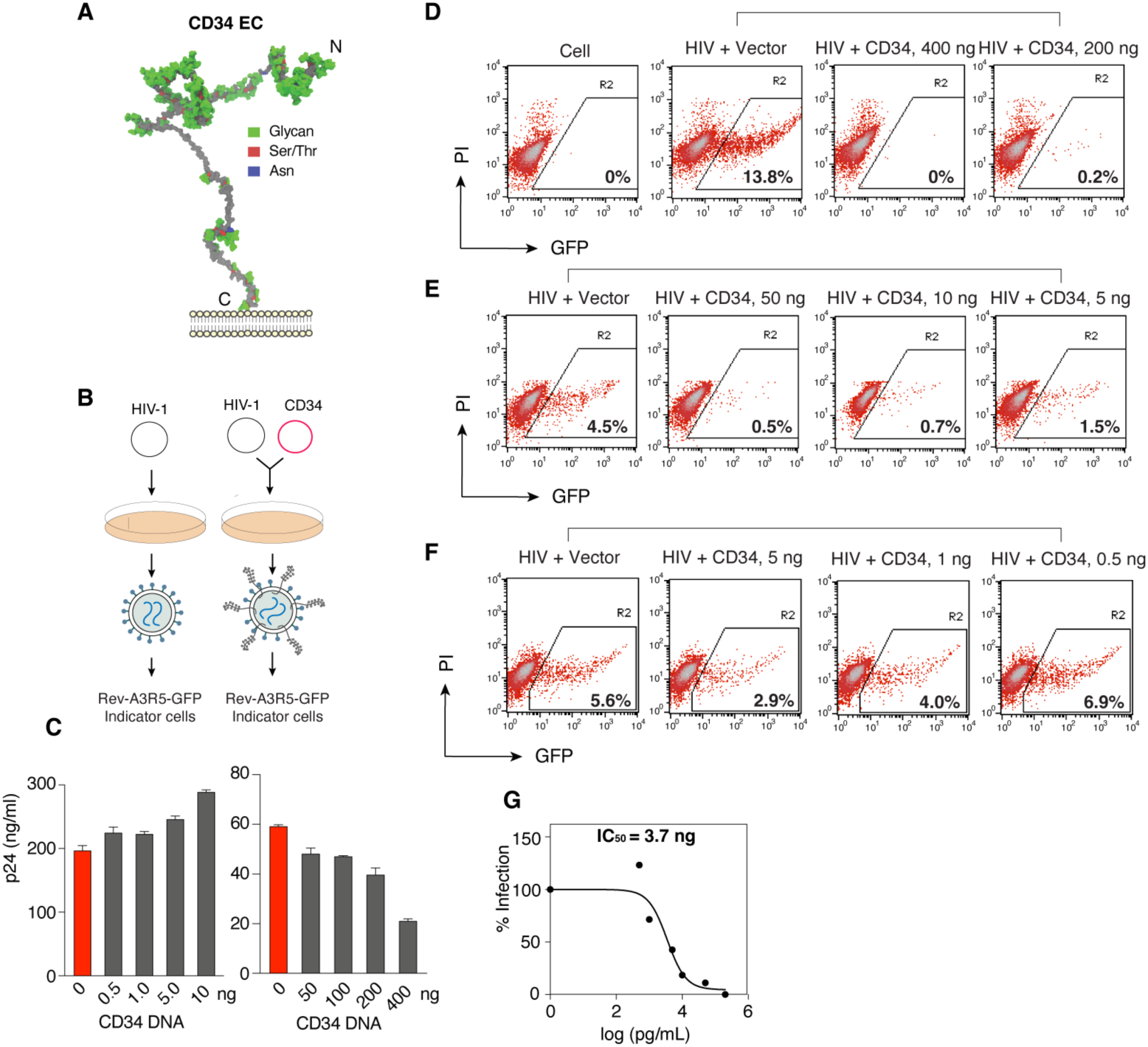
Dosage-dependent inactivation of HIV-1 virion infectivity by CD34. (**A**) Structure of the CD34 extracellular domain produced from Rosetta. The protein chain is shown in grey, with Asn in blue, Ser/Thr in red, and glycans in green. (**B**) Schematic of virion production in the presence of CD34 in virus producer cells. (**C**) Effects of CD34 on HIV-1 virion release. HEK293T cells were co-transfected with HIV-1 DNA (1 μg) plus a CD34-expression vector (0.5 to 400 ng of DNA). Virion release was quantified at 48 h post co-transfection by HIV-1 p24 ELISA. Data are represented as mean ± SD from assay triplicate. (**D** to **F**) CD34 inactivates HIV-1 virion infectivity. HEK293T cells were co-transfected with HIV-1 DNA (1 μg) plus a CD34-expressing vector (0.5 to 400 ng of DNA). Virions were harvested at 48 h and viral infectivity was quantified by infecting Rev-A3R5-GFP indicator cells using an equal level p24 viral input. Shown are the percentages of GFP^+^ cells at 48 h post infection. The experiment was repeated more than three times. In all co-transfection experiments, an empty vector (Vector) was used to ensure that an equal amount of total plasmid DNA was used in all co-transfection. (**G**) The 50% inhibition dosage (IC_50_) of CD34 was calculated based on the averages of infectivity assays.

To investigate possible antiviral properties of CD34 on stem/progenitor cells, we first assembled HIV-1 particles in the presence of CD34. HEK293T virus-producer cells were co-transfected with a CD34-expressing vector and HIV-1 DNA to produce viruses (**Figure 1B**). We first examined whether expression of CD34 in producer cells inhibits virion release, and tested a range of CD34 vector dosages, from 0.5 ng to 400 ng. We found that at low dosages (0.5–10 ng), CD34 did not inhibit virion release. At higher dosages, CD34 only slightly inhibited virion release (approximately 50% inhibition at 400 ng) (**Figure 1C**). We further quantified the infectivity of progeny virions using the HIV Rev-dependent GFP indicator cell, Rev-A3R5-GFP ^27,47,48^. When an equal p24 level of input virus was used, we observed near-complete inhibition of HIV virion infectivity at CD34 dosages of 10 ng and above. CD34 partially inhibited HIV-1 infectivity at inputs as low as 1 ng (CD34-to-HIV DNA ratio, 1:1000), and there was a dose-dependent inactivation of HIV-1 at CD34 vector doses from 1 to 10 ng (**Figure 1D** to **1F**). The half-maximal inhibitory dosage (IC_50_) was quantified to be at 3.7 ng of CD34 (**Figure 1G**).

CD34 is a sialomucin, and shares the same selectin ligands as PSGL-1 ^39,40^. Therefore, we hypothesized that mechanistically, CD34-mediated inhibition of virion infectivity may result from virion incorporation that either sterically hinders virus particle attachment to target cells and/or blocks HIV Env virion incorporation, mechanisms similar to those of PSGL-1 ^27,28^. To determine virion incorporation of CD34, we performed density gradient centrifugation to purify virions released from co-transfected HEK293T cells and observed co-sedimentation of CD34 with purified virions only when cells were co-transfected with the CD34 vector plus HIV DNA (**Figure 2A** and **2B**), but not with the CD34 vector alone (**Figure 2C**). We further confirmed CD34 virion incorporation using a particle pull-down assay ^41^; anti-CD34 antibody-conjugated magnetic beads were used to pull down virion particles that express CD34 on the surface (**Figure 2D**). Magnetically purified virions were further quantified for the presence of HIV-1 p24 in the virions. We were able to demonstrate that only the anti-CD34 antibody, but not the control non-specific antibody, selectively pulled down p24^+^ virion particles in a CD34-dosage dependent manner, confirming virion incorporation of CD34 (**Figure 2E**). We also investigated the effects of CD34 on virion Env incorporation, but did not observe inhibition of Env incorporation at the CD34 dosages tested (**Figure 2F**). This is different from PSGL-1, which blocks Env incorporation ^27^. We further performed a virion attachment assay and observed that CD34-imprinted virions were inhibited in their ability to attach to Rev-A3R5-GFP target cells (**Figure 2G**). To determine whether this inhibition of virion attachment is dependent on specific interaction of HIV Env with the CD4 receptor and co-receptor, we performed a virion attachment assay using CD4^+^ and CD4^-^ cells (HeLa JC.53 and HeLa), and observed that the presence of CD34 inhibited virion attachment to both (**Figure 2H**), demonstrating that the inhibition of virion attachment is not dependent on interaction of Env with the CD4 receptor. In addition, we took advantage of previous findings that HIV particles can bind target cells even in the absence of Env-receptor interaction ^49,50^. We assembled HIV particles devoid of any viral envelope glycoproteins (NL4-3/KFS) ^51^ in the presence or absence of CD34, and performed virion attachment assays. As shown in **Figure 2H**, we observed that CD34-imprinted, Env-negative particles were also impaired in their ability to attach to HeLa and HeLa JC.53 cells. These results confirmed that CD34 inhibits virion attachment to cells in an envelope glycoprotein-independent manner, a phenotype similar to that of PSGL-1. Based on these results, we speculate that the anti-viral activity of CD34 may be pleotropic, and thus we further tested the antiviral activities of CD34 against an ectotropic retrovirus, murine leukemia virus (MLV-GFP); CD34 was found to also inhibit replication of MLV (**Figure S1)**.

**Figure 2.**
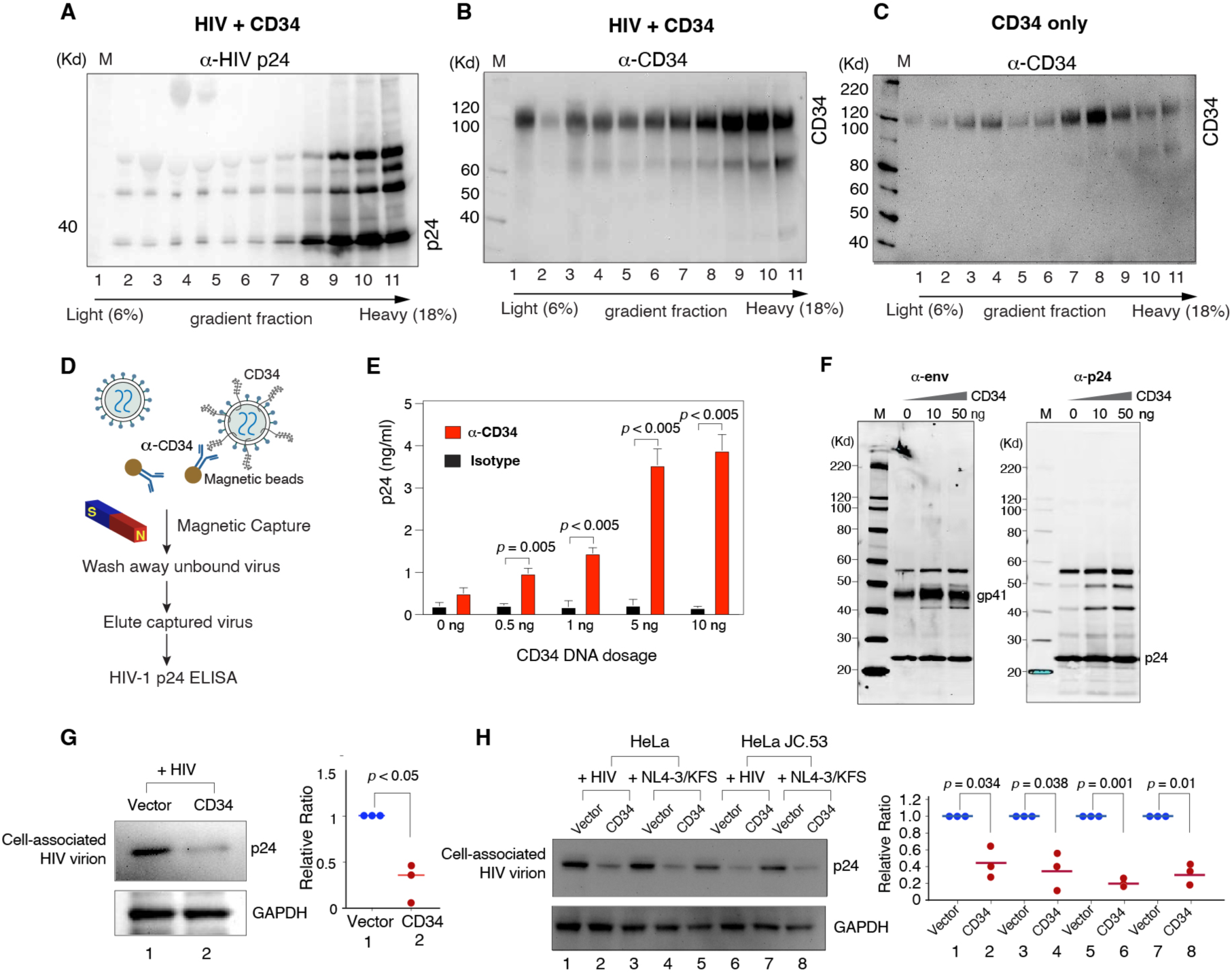
Virion incorporation of CD34 inhibits virus attachment to target cells. (**A** to **C**) Virion incorporation of CD34. HEK293T cells were co-transfected with 10 μg HIV-1 DNA plus 2 μg of CD34 DNA (HIV + CD34) or with CD34 DNA only (CD34 only). Supernatants were harvested at 48 h and purified through 6%–18% OptiPrep gradient ultra-centrifugation. CD34 and viral p24 proteins were analyzed by western blot using antibodies against HIV-1 p24 or CD34. Shown is the representative of three repeats. (**D** and **E**) Schematic of the immunomagnetic capture assay for detecting CD34 on HIV-1 particles (**D**). HEK293T cells were co-transfected with HIV-1 DNA (1 μg) plus a CD34 expression vector (0.5–10 ng) or an empty vector. Supernatants were harvested at 48 h and incubated with magnetic beads coated with anti-CD34 antibody or an isotype control antibody. Captured particles were washed, eluted, and quantified for p24. *p*-values were based on three independent assays (**E**). (**F**) Effects of CD34 on HIV-1 Env incorporation. HEK293T cells were co-transfected with HIV-1 DNA plus a CD34 expression vector (10 and 50 ng) or an empty vector. Particles were harvested at 48 h and purified through a sucrose gradient. Virions were lysed and analyzed with western blot using anti-HIV-1 gp41 or p24 antibodies. Representative blots from three independent repeats are shown. (**G**) CD34 blocks HIV-1 virion attachment to target cells. Viral particles were produced by co-transfection of HEK293T cells with HIV-1 DNA (1 μg) and a CD34 expression vector or an empty vector (10 ng). HIV-1 p24-normalized viral particles were then assayed for attachment to target Rev-A3R5-GFP cells by western blot for cell-bound p24. Representative blots from three independent assays are shown. The protein band intensity was quantified and the relative ratios of vector/GAPDH and CD34/GAPDH were calculated. Vector/GAPDH ratios were assigned as “1”. (**H**) CD34 blocks HIV-1 virion attachment independent of Env interaction with cellular receptors. Viral particles were produced similarly by co-transfection of HEK293T cells with HIV-1 DNA or HIV-1(NL4-3/KFS) and a CD34 expression vector or an empty vector. HIV-1 p24-normalized viral particles were then assayed for attachment to target HeLa or HeLa JC.53 cells by western blot for cell-bound p24. Representative blots from three independent assays are shown. The protein band intensity was similarly quantified. All *p*-values were calculated using a two-tailed T-test.

### Expression of CD34 in Human CD4 T Cells Elicits Anti-HIV-1 Activity

To confirm that CD34 has antiviral activity in HIV-1 target CD4 T cells, we transiently expressed CD34 in a human CD4^+^ lymphoblastoid cell line, Rev-A3R5-GFP ^47,52^ (**Figure 3A**). Following CD34 vector electroporation, surface expression of CD34 on Rev-A3R5-GFP was confirmed by surface staining (**Figure 3B**). Cells were subsequently infected with HIV-1, and viral replication was monitored by HIV-1-mediated GFP expression. We observed that expression of CD34 on target cells inhibited HIV-1 replication in CD34^+^CD4^+^ T cells, reducing the percentage of HIV-infected GFP^+^ CD4 T cells from 35.6% to 8.4% (a 76.4% reduction) (**Figure 3C**). The experiments were repeated three times, and on average, there was a statistically significant 72.6% reduction of HIV replication in CD34^+^CD4^+^ T cells. Given that CD34 does not block viral release but inactivates progeny virion infectivity (**Figure 1**), this inhibition likely results from blocking secondary infection. We further mapped the effects of CD34 on HIV infection by using an HIV-1 Env-pseudotyped, single-cycle virus, HIV(KFS)(Env) ^51,53^. Expression of CD34 on target cells led to a 34.6% reduction of the single-cycle viral infection (from 2.6% to 1.7%) (**Figure 3D**). We repeated the experiment three times, and on average, there was a statistically significant but modest 38.9% reduction in HIV(KFS)(Env) infection. This slight reduction of single-round infection likely resulted from inhibition of viral entry. We performed beta-lactamase-based viral entry assays (4 independent assays) and found that on average, expression of CD34 slightly reduced viral entry from 64.9% to 55% (a 15% reduction) (**Figure 3E**). Collectively, these results demonstrate that CD34 expression on target cells only modestly affected viral entry and release (**Figure 3G** and **1C**), whereas progeny virion incorporation of CD34 potently blocked particle attachment to target cells (**Figure 2G** and **2H**). Therefore, we conclude that CD34 inhibits HIV-1 infection mainly by blocking the secondary but not the primary infection through inactivating progeny virions released.

**Figure 3.**
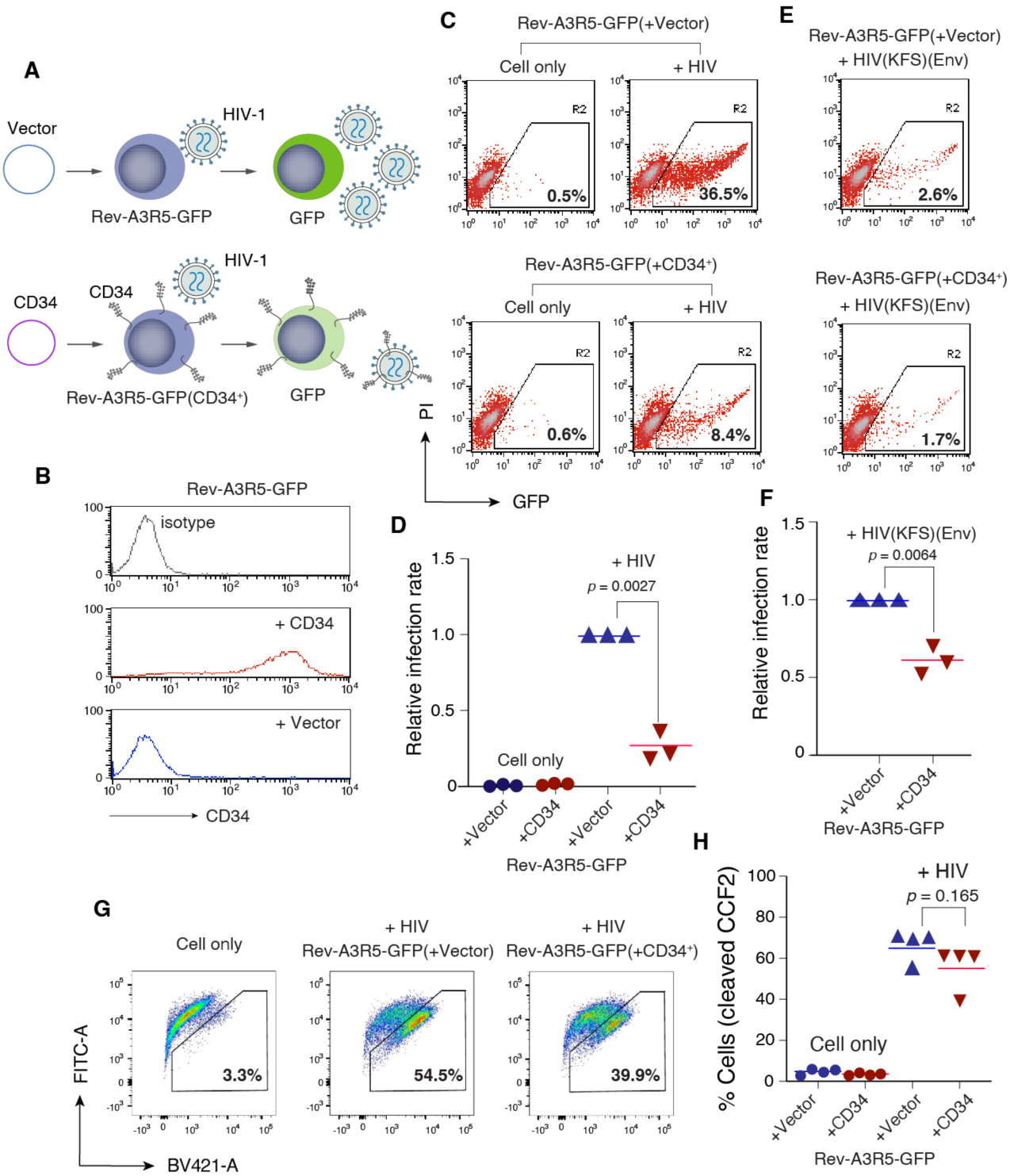
Expression of CD34 in human CD4 T cells inhibits HIV infection. (**A**) Schematic of the assay to determine effects of CD34 expression on human CD4 T cells. (**B**) Quantification of surface CD34 expression on a human CD4 T cell, Rev-A3R5-GFP, following electroporation of cells with a CD34 expression vector or a control empty vector. (**C** and **D**) Expression of CD34 in Rev-A3R5-GFP T cells inhibits HIV infection. Cells were electroporated with a CD34-expression vector or a control empty vector. At 48 h post electroporation, cells were infected with HIV-1, and Rev-dependent expression of GFP was quantified at 3 days post infection (**C**). The experiments were independently repeated three times. For statistical analysis, the infection rates (% GFP^+^ cells) of empty vector-electroporated cells were assigned as “1”. (**E** and **F**) Cells were also similarly electroporated with DNA and then infected with a single-cycle virus, HIV-1(KFS)(Env). HIV-1 infection was quantified by GFP expression at 3 days post infection. (**G**) Effects of CD34 on HIV-1 viral entry. Rev-A3R5-GFP CD 4 T cells were electroporated with a CD34-expression vector or a control empty vector. At 48 h post electroporation, cells were infected with HIV-1 for 2 hours for viral entry assay (BlaM) using an equal level p24 virus input. The percentages of cells with cleaved CCF2 are shown. The experiments were independently repeated 4 times. All *p*-values were calculated using a two-tailed T-test.

### Physiological Levels of CD34 Naturally Expressed on CD34^+^ Hematopoietic Stem And Progenitor Cells Are Functional in Limiting Retroviral Replication

To confirm that CD34 naturally expressed on stem and progenitor cells possesses antiviral activity, we examined HIV infection of primary human CD34^+^ cord blood hematopoietic stem and progenitor cells (HSPC) that express CXCR4 but not the CD4 receptor of HIV-1 (**Figure 4A**). For infection, we used a VSV-G (vesicular stomatitis virus glycoprotein) pseudotyped HIV-1, HIV-1(VSV-G), which bears both HIV Env and VSV-G on the surface (**Figure 4B**); VSV-G pseudotyping allowed HIV CD4-independent entry and single-round infection of the CD34^+^ cord blood cells. HIV-1 particles released from this single-round infection contain HIV-1 Env and CD34, and thus can be quantified for virion infectivity using Rev-A3R5-GFP (**Figure 4C**). As shown in **Figure 4D**, direct infection of CD34^+^ cord blood cells allowed HIV-1(VSV-G) entry and low-levels of virion release **(Figure 4D)**. However, when the infectivity of the progeny virions was quantified on Rev-A3R5-GFP, no or minimal infectivity was detected (**Figure 4E** and **Figure S2**). To further probe the role of CD34, we took advantage of a previous finding that culturing hematopoietic progenitor cells (HPCs) in GM-CSF and TNF-α can induce downregulation of CD34 from cell surface ^8^. We cultured CD34^+^ cord blood HSPCs in either the stem cell expansion medium (ExpM) alone or ExpM plus GM-CSF/TNF-α for 7 days.

**Figure 4.**
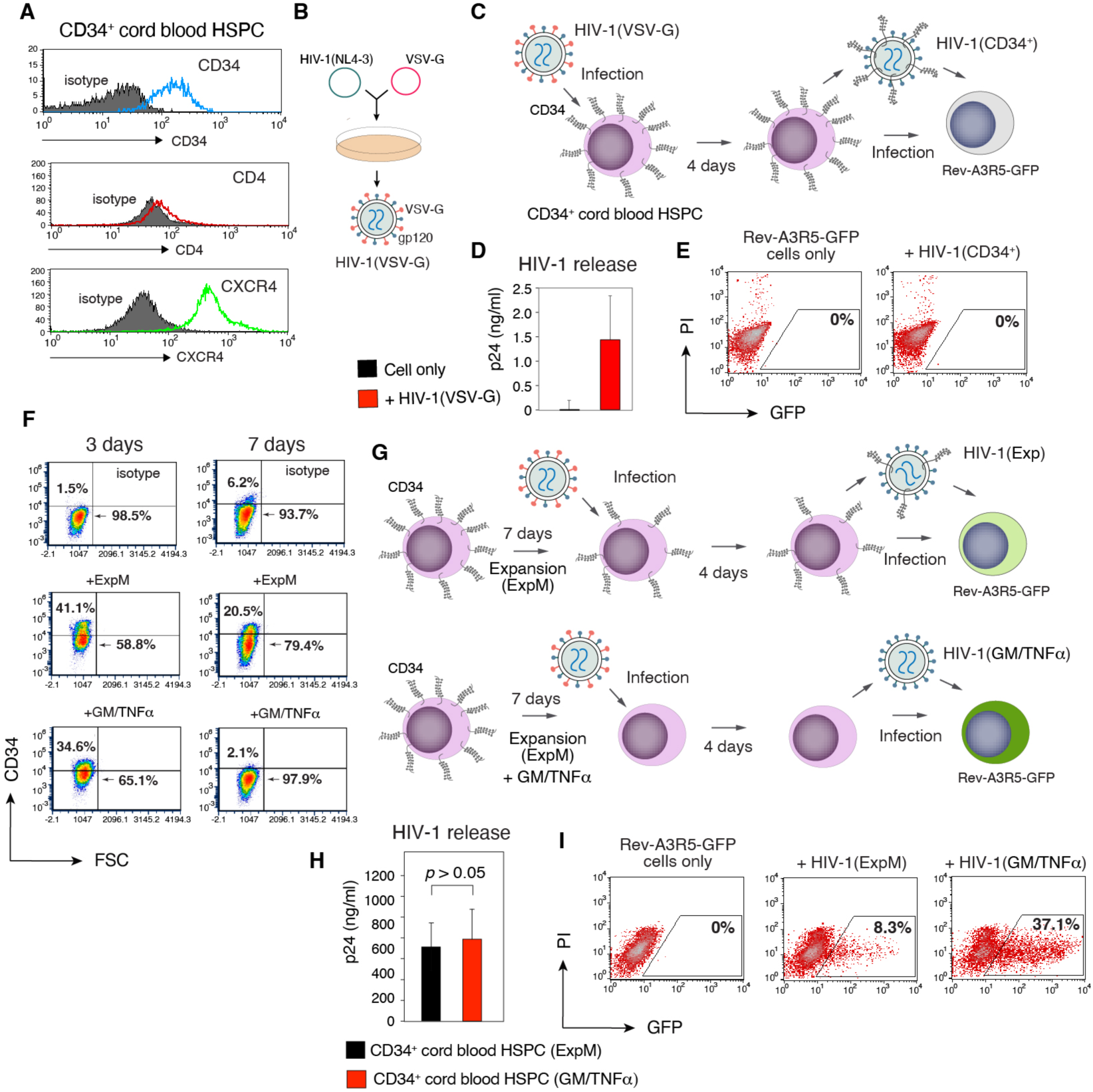
GM-CSF/TNF-α-mediated downregulation of CD34 enhances the infectivity of HIV progeny virions released from CD34^+^ cord blood hematopoietic stem/progenitor cells (HSPCs). (**A**) Measurement of surface CD34, CD4, and CXCR4 expression on CD34^+^ cord blood HSPCs. (**B**) Schematic of the assembly of HIV-1(VSV-G). (**C** to **E**) Direct infection of CD34^+^ cord blood HSPCs led to low-level release of non-infectious HIV particles. Schematic of the infection procedures to determine HIV particle release and virion infectivity (**C**). CD34^+^ cord blood HSPCs were directly infected with HIV-1(VSV-G) for 4 days, and virion release was quantified with p24 ELISA (**D**). Virions released were also quantified for infectivity by infecting Rev-A3R5-GFP indicator cells, and GFP expression was quantified at 8 days post infection. (**F**) GM-CSF/TNF-α-mediated downregulation of CD34 on CD34^+^ cord blood HSPCs. Cells were cultured in stem cell expansion medium (ExpM) alone or ExpM plus GM-CSF/TNF-α for 7 days, and surface expression of CD34 was measured. (**G** to **I**) Schematic of the infectivity assay to demonstrate enhancement of HIV virion infectivity by GM-CSF/TNF-α (**G**). CD34^+^ cord blood HSPCs were cultured in ExpM or in ExpM plus GM-CSF/TNF-α for 7 days. Cells were infected with HIV-1(VSV-G). Following infection for 4 days, virion release was quantified with HIV p24 ELISA (**H**). Virions released were also quantified for infectivity by infecting Rev-A3R5-GFP indicator cells using an equal level p24 viral input (**I**).

Culturing cells in ExpM partially downregulated CD34, whereas culturing cells in GM-CSF/TNF-α completely downregulated CD34 (**Figure 4F**). We subsequently infected cells with HIV-1(VSV-G), and the progeny virions were harvested and quantified (**Figure 4G**). As shown in **Figure 4H**, GM-CSF/TNF-α culturing did not enhance viral release. However, when the infectivity of the virions released was quantified on Rev-A3R5-GFP, using an equal p24 level of input virus, we observed a GM-CSF/TNF-α-mediated enhancement of virion infectivity (**Figure 4I** and **Figure S3**) that correlates with GM-CSF/TNF-α-mediated CD34 downregulation in the virus-producer cells.

To recapitulate the observation and for more mechanistic studies, we repeated the experiment using an undifferentiated leukemia CD34^+^ progenitor cell, Kasumi-3 ^54^, that naturally expresses the HIV receptor and co-receptor, both CD4 and CXCR4 (**Figure 5A** and **5B**, and **Figure S4**). This allowed us to directly use HIV-1 without relying on pseudotyping. In addition, the levels of CD34 on Kasumi-3 were comparable to those expressed on primary human bone marrow CD34^+^ cells (**Figure 5A**). We directly infected or transfected Kasumi-3 cells with HIV-1 virus or viral DNA, both HIV-1(NL4-3) and HIV-1(89.6), and detected low-levels of viral release; however, there was minimal spreading viral replication (**Figure 5C** and **Figure S5**), in agreement with a previous finding showing that HIV-1 can enter and express genes in CD34^+^ multipotent hematopoietic progenitor cells (HPCs) ^8^. To confirm the role of CD34 in HIV-1 spreading infection of Kasumi-3 cells, we similarly cultured Kasumi-3 in GM-CSF/TNF-α, which also induced the downregulation of CD34, as seen in the CD34^+^ cord blood HSPC (**Figure 5D** and **5E**, and **Figure S6**). To determine the effects of GM-CSF/TNF-α-induced CD34 downregulation on the infectivity of progeny virions released, we electroporated HIV-1 DNA into cells to assemble virion particles (**Figure 5F**); we used electroporation for virus assembly to minimize CD34 effects on HIV entry (**Figure 3G**). As a control, viral particles were also assembled in the absence of GM-CSF/TNF-α. Culturing Kasumi-3 cells in GM-CSF/TNF-α did not enhance viral release (**Figure 5G**). However, when the infectivity of the virions released was quantified using an equal p24 level of input virus, we observed enhanced infectivity of virions released from GM-CSF/TNF-α-cultured Kasumi-3 cells (**Figure 5H**), a phenotype recapitulating what we have observed in GM-CSF/TNF-α-treated CD34^+^ cord blood HSPC (**Figure 4G** to **4I**).

**Figure 5.**
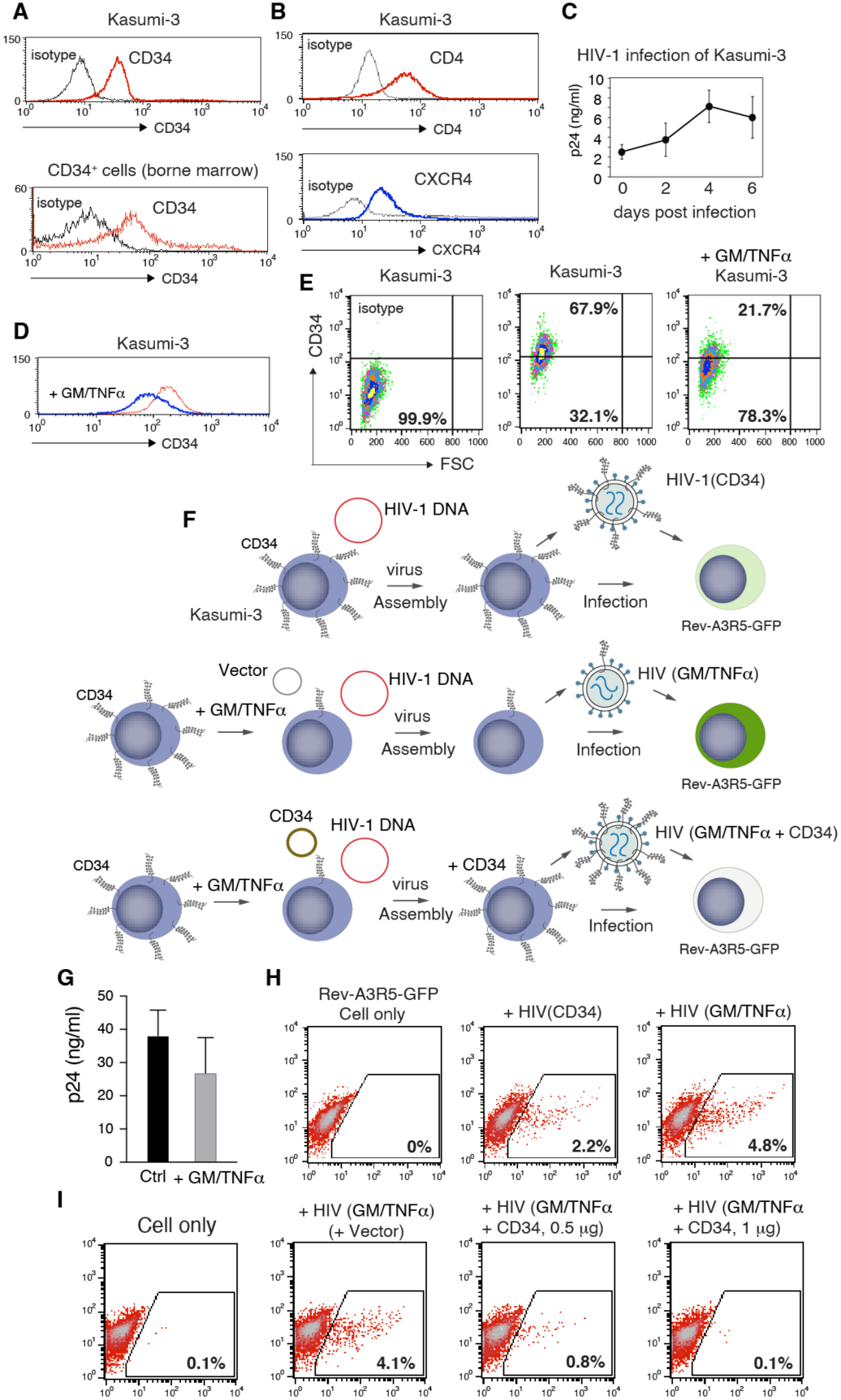
GM-CSF/TNF-α-mediated downregulation of CD34 enhances the infectivity of HIV progeny virions released from CD34^+^ Kasumi-3 cells. (**A**) Measurement of surface CD34 expression on Kasumi-3 cells and on primary human bone marrow CD34^+^ cells. (**B**) Measurement of surface CD4 and CXCR4 expression on Kasumi-3 cells. (**C**) Direct HIV-1 infection of Kasumi-3 cells. Cells were infected with HIV-1(89.6) for 4 h, washed, and then cultured for 6 days. Viral p24 release was quantified by ELISA. The experiments were repeated 3 times. (**D** and **E**) Downregulation of CD34 by GM-CSF/TNF-α. Cells were cultured in GM-CSF/TNF-α or medium for 7 days, and surface expression of CD34 was measured. (**F** to **I**) Schematic of the infectivity assays to demonstrate enhancement of HIV virion infectivity by GM-CSF/TNF-α. Kasumi-3 cells were cultured in GM-CSF/TNF-α or medium for 7 days, and then electroporated with HIV-1 DNA or with HIV-1 DNA plus a CD34-expression vector or an empty vector for control. Viral particles were harvested at 3 days post electroporation, and virus infectivity was quantified by infecting Rev-A3R5-GFP indicator cells using equal p24 levels of input viruses.

To determine whether downregulation of CD34 by GM-CSF/TNF-α contributes to GM-CSF/TNF-α-mediated enhancement of virion infectivity, we reintroduced CD34 into GM-CSF/TNF-α-cultured Kasumi-3 cells using a CD34 expression vector with constitutive CMV promoter (**Figure 5F**). Over-expression of CD34 in GM-CSF/TNF-α-cultured Kasumi-3 virus producer cells diminished HIV infectivity enhanced by GM-CSF/TNF-α (**Figure 5I**). Although it is possible that GM-CSF/TNF-α may also downregulate other membrane proteins, our results at a minimum suggest that CD34 is one of the factors responsible for GM-CSF/TNF-α-mediated enhancement of virion infectivity.

To conclusively define the role of CD34 in limiting HIV replication in CD34^+^ stem/progenitor cells, we further performed CRISPR/Cas9 knockout of CD34 in Kasumi-3 cells. The disruption of the CD34 gene was confirmed by DNA sequencing, and by western blot quantification and cell surface staining of CD34 (**Figure 6A** to **6C**). CD34*^ko^*Kasumi-3 cells were subsequently infected with an HIV-1(GFP) reporter virus, HIV-1(NLNEG1-ES-IRES) ^55^, and we observed enhancement of HIV-1 replication in CD34*^ko^*Kasumi-3 cells in comparison with the CD34^+^ parental Kasumi-3 cells (**Figure 6D** and **Figure S7**). To further confirm that the enhancement of HIV-1 replication in CD34*^ko^*Kasumi-3 was indeed derived from enhanced virion infectivity, we harvested progeny virus particles 4 days after infection and quantified HIV infectivity on Rev-A3R5-GFP reporter cells using an equal p24 viral input. As shown in **Figure 6E**, HIV-1 particles released from CD34*^ko^*Kasumi-3 cells have a much greater infectivity compared with those released from the parental CD34^+^ Kasumi-3 cells (39.1% versus 0.1%) (**Figure S8**). We further confirmed the results using wild-type HIV-1 infection of the CD34*^ko^*Kasumi-3 cells. Again, we observed enhanced HIV replication in the CD34 knockout cells (**Figure 6F** and **Figure S9**). Collectively, our results conclusively demonstrated that CD34 expressed naturally on CD34^+^ hematopoietic stem/progenitor cells is functional in diminishing the infectivity of virions released from infection, limiting HIV spreading infection among CD34^+^ stem/progenitor cells.

**Figure 6.**
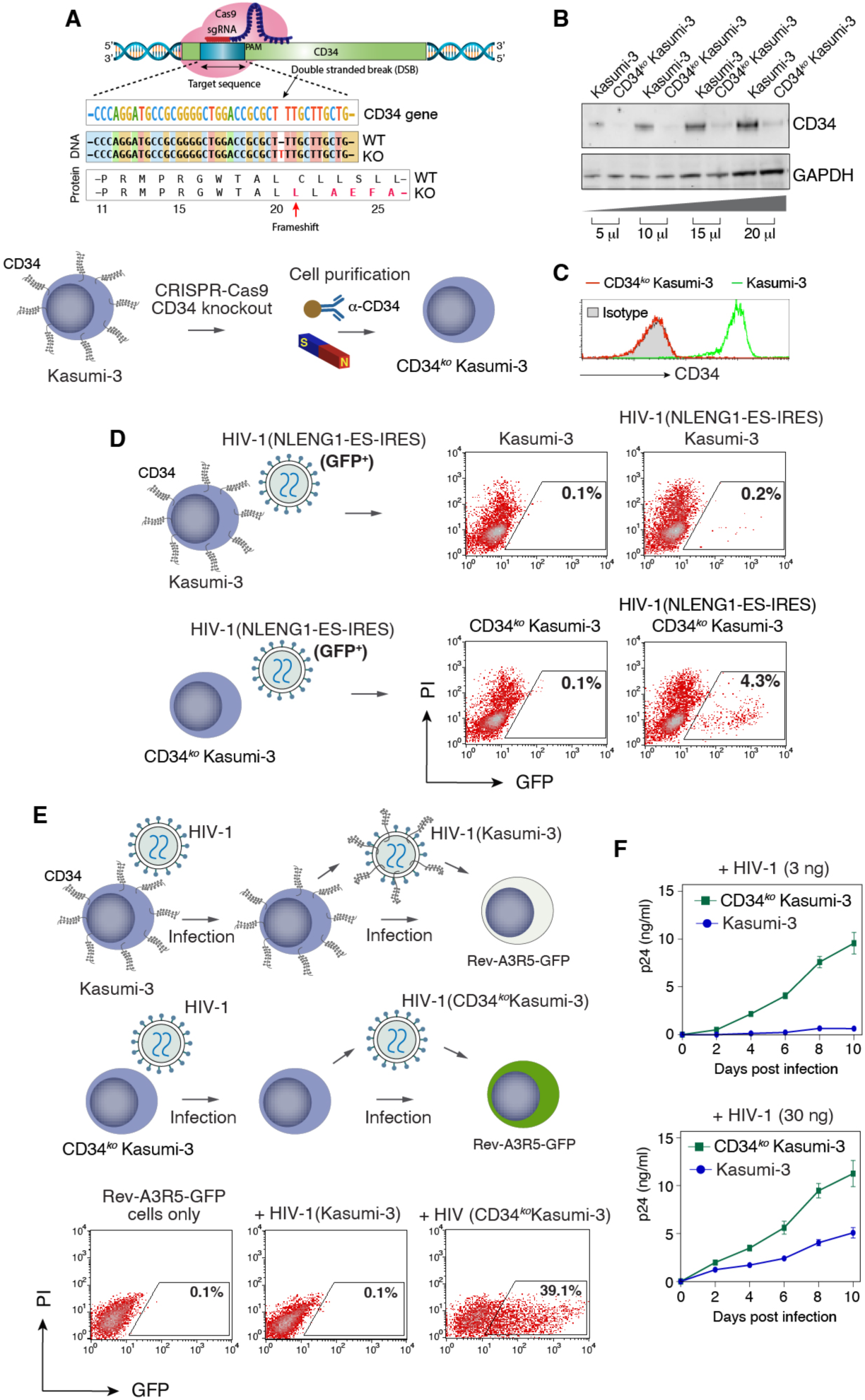
CRISPR/Cas9 knockout of CD34 leads to enhanced HIV virion infectivity. (**A**) Schematic of the CRISPR/Cas9 knockout of CD34 on Kasumi-3 and the cell purification procedures. The disruption of the CD34 gene was confirmed by DNA sequencing showing an insertion of “T” which causes a protein frame shift. (**B** and **C**) Confirmation of CD34 knockout by western blot (**B**) and cell surface staining of CD34 and flow cytometry (**C**). (**D**) Enhanced HIV-1 replication in CD34^ko^ Kasumi-3 cells. Kasumi-3 or CD34^ko^ Kasumi-3 cells were identically infected with HIV-1(NLENG1-ES-IRES) using an equal p24 viral input. At 2 days post infection, viral replication was quantified by measuring GFP expression. Shown are results from three independent experimental replicates. (**E**) Enhanced infectivity of HIV-1 virions released from CD34^ko^ Kasumi-3 cells. Kasumi-3 or CD34^ko^ Kasumi-3 cells were infected with HIV-1 virus. At 4 days post infection, virions released were harvested and virion infectivity was quantified by infecting Rev-A3R5-GFP indicator cells using an equal p24 viral input. The percentage of GFP^+^ cells was quantified at 10 days post infection by flow cytometry. Shown are results from three independent experimental replicates. (**F**) CRISPR/Cas9 knockout of CD34 leads to enhanced HIV spreading infection in CD34^ko^ Kasumi-3 cells. Kasumi-3 and CD34^ko^ Kasumi-3 cells were identically infected with HIV-1 using an equal p24 viral input. HIV replication was quantified by measuring HIV p24 levels in the culture supernatant.

## Discussion

In this article, we demonstrated that the ubiquitous stem cell marker CD34 is an antiviral protein limiting retroviral spreading infection and proliferation among CD34^+^ stem and progenitor cells. It has long been recognized that direct HIV-1 infection of CD34^+^ stem cells and progenitor cells is difficult and limited ^5,8,13,17,56^. Some early studies failed to detect HIV-1 DNA in a highly purified CD34^+^ cell population ^57^ or in Lin(-)/CD34(+) hematopoietic precursor cells in patients ^58^, while others have suggested that CD133^+^ hematopoietic progenitor cells harbor HIV genomes in a subset of patients on long-term ART ^59^. HIV has also been found to directly infect CD34^+^ multipotent hematopoietic progenitor cells (HPCs), although viral replication is limited ^8,14^. Importantly, recent studies have suggested that CD34^+^ hematopoietic stem cells and progenitor cells can serve as a distinct HIV reservoir that contributes to HIV persistence in ART-suppressed patients ^10,60^. Our detailed mechanistic studies of the role of CD34 demonstrated that: (1) CD34 does not intrinsically block viral entry and release, allowing viral entry, integration, and gene expression to occur in CD34^+^ cells; (2) a dominant phenotype of CD34 is the inactivation of progeny virions released, limiting spreading viral replication in the CD34^+^ cell population; (3) our study also supports a unique potential of HIV^+^ CD34^+^ cells for producing infectious virus following cell differentiation and the loss of surface CD34 ^9,16,60^.

### Why Do Stem Cells Utilize CD34 For Protection While They Are Protected by Restriction Factors And Intrinsic Interferon Stimulated Genes (ISGs)?

Previous studies have demonstrated that stem cells are highly resistant to viral infection compared to their differentiated progeny ^2-6^, and this resistance is linked to the stem cell-specific expression of restriction factors (*e.g.*, ZFP809) and intrinsic interferon stimulated genes (ISGs) (*e.g.*, IFITM1) that are constitutive and refractory to interferon induction ^3,7^. It has also been shown that the expression of these intrinsic ISGs is selective and cell-type specific rather than in full spectrum ^7^. For instance, intrinsic ISGs with anti-proliferative properties might not be present in self-renewing stem cells ^7,61^. As such, there may be cases where effective intrinsic ISGs are absent or are antagonized by viruses, presenting the need for additional layers of protection. Most intrinsic ISGs function by blocking early steps in viral infection. In contrast, CD34 does not prevent viral entry or early steps but rather inactivates progeny virions, inhibiting the spread of viruses among stem cells (**Fig. 1** and **Fig. 2**).

Functionally, the antiviral properties of CD34 complement those of intrinsic ISGs, targeting viruses at different stages of their life cycle. Interestingly, CD34 also share similarities with intrinsic ISGs, as their expressions are constitutive in stem cells and decline upon stem cell differentiation (**Fig. 4**) ^7^. The loss of CD34 and intrinsic ISGs following differentiation may be beneficial, allowing the onset of interferon-inducible ISGs, which have broader spectrum and higher levels of specificity and regulation. This transition could enhance antiviral efficiency while minimizing the potential negative effects of ISGs ^7^.

### Why Does CD34 Inactivate Retrovirus Progeny Virion Rather Than Block Viral Entry into Stem Cells?

While widespread infection of retroviruses in stem cells can be detrimental, limited retroviral infection and genome integration should be tolerable and potentially important for shaping the innate immune landscape through virus-host interactions ^7^. For instance, the endogenous retrovirus (ERVs) HERVK(HML-2) contains an accessory protein Rec, and its overexpression in pluripotent cells elevates IFITM1 expression, which helps inhibit viral infection ^62^. Remarkably, HERVK viral-like particles (VLPs) and the Gag protein have also been detected in human blastocysts, indicating the presence of retroviral components during early human development ^62^. The human genome harbors a substantial collection of endogenous retroviruses, constituting about 8% of the genome, which are remnants from ancient retroviral infections ^63^. Thus, it may not be evolutionarily advantageous or feasible to completely block retroviral entry ^64^. It’s also tempting to speculate that the presence of CD34 on virion particles would prevent their re-entry and spread in stem cells in the event of a viral breakthrough and particle release. In this context, CD34 could act as a “guardrail”, compelling retroviruses to co-evolve within cells while managing the risk of full-scale infection.

### Why Cannot HIV Accessory Proteins Downregulate CD34 While They Antagonize Restriction Factors and ISGs?

HIV has evolved a range of accessory proteins such as Vif, Vpu, and Nef to antagonize ISGs ^18-28^. Notably, while HIV can use accessory proteins such as Vpu and Nef to antagonize mucin-like proteins such as PSGL-1 in human CD4 T cells ^26,27^, the virus is not capable of downregulating CD34 (**Fig. S10**). This suggests that there is an inherent protective mechanism in stem and progenitor cells that limits retroviral spread in these crucial cell types.

The role of stem cells is vital in development, self-renewal, and tissue regeneration, indicating that they likely possess multiple layers of defense against viral infection. The first line of protection comes from the expression of intrinsic ISGs ^7^, while CD34 appears to serve as a final barrier. The evolution of HIV and its accessory proteins points to a complex interplay between the virus and host defenses, allowing HIV to break through some ISGs to integrate and establish a reservoir within a subset of human stem cells. Indeed, CD34^+^ multipotent hematopoietic progenitor cells have been found to harbor HIV proviral DNA, which underscores the virus’s ability to bypass certain ISGs for integration and establishing a reservoir ^8,10,14,60^. However, the presence of CD34 on stem cells implies that the production of infectious virus can only happen after these cells differentiate and lose CD34 expression. This positions CD34 as a critical “guardrail” against HIV, suggesting that stem cells may have yet-to-be-discovered mechanisms to prevent early downregulation of CD34 by viral accessory proteins.

In summary, our study underscores the role of CD34 as a crucial new host factor in stem cell immunity and a significant contributor to our understanding of retrovirus-host interactions in evolution. Given the essential role of stem cells in development, self-renewal, and tissue regeneration, they possess multiple layers of defense against viral infections. While restriction factors and intrinsic ISGs serve as the first line of defense by blocking early viral steps, CD34 acts as a final barrier that can neutralize progeny viruses released from cells, even if these viruses manage to overcome ISG restrictions. The human genome contains a large pool of endogenous retroviruses. With stem cells undergoing high proliferation and active retroviral gene expression during early development, additional protection is critical, especially against complex retroviruses like HIV, which have evolved accessory proteins to escape ISG defenses. CD34 provides an essential second layer of protection beyond ISGs. Importantly, unlike ISGs, CD34 is resistant to viral downregulation, thereby functioning as a final “guardrail” that compels retroviruses to co-evolve within the cells while mitigating the risk of widespread infection.

## ACKNOWLEDGEMENTS

We wish to thank the NIH AIDS Reagent Program for reagents, Jennifer Guernsey for editing assistance. This work was supported by National Institutes of Health grant R01AI148012 (YW) and National Institutes of Health grant R56AI183995 (YW, DK, and MSJ)

## AUTHOR CONTRIBUTIONS

Conceptualization, Y.W.; methodology, YW, SH, AH, ST, DD, BH, YH, LS, and CL; formal analysis, YW, SH, AH, ST, DD, LS, and CL; investigation, YW, SH, AH, ST, DD, JLY, BH, YH, CL; writing – original draft, Y.W.; writing – review and editing, YW, SH, AH, ST, DD, LS; supervision, Y.W.; funding acquisition, Y.W.

## SUPPLEMENTAL INFORMATION

Supplemental **Figures S1** to **S10**

## METHODS

### Cells and Cell lines

Anonymous human cord blood samples were collected at the Cleveland Cord Blood Center. EasySep™ Human Cord Blood CD34 Positive Selection Kit III (Stem Cell Technologies) was used to purify CD34+ cells from cord blood. Purified CD34+ cord blood hematopoietic stem and progenitor cells (HSPCs) were cultured in StemSpan SFEM II medium (Stemcell) with or without expansion media (ExpM, cytokine mixture of SCF, Flt3L, and TPO at 100 ng/ml) (Stemcell). CD34+ cord blood HSPCs were also cultured in ExpM with or without recombinant human GM-CSF (R&D Systems) (100 ng/ml) plus recombinant human TNF-α (Biolegend) (2.5 ng/ml) for 7 days. CD34+ Kasumi-3 (ATCC) cells were maintained in RPMI-1640 plus 20% FBS supplemented with 100 units/ml of penicillin and 100 µg/ml of streptomycin (Invitrogen) and 50 µg/ml of gentamicin (Sigma-Aldrich). HEK293T (ATCC) and NIH/3T3 (ATCC) cells were maintained in Dulbecco’s modified Eagle’s medium (DMEM) (Invitrogen) containing 10% heat-inactivated FBS and 100 units/ml of penicillin and 100 µg/ml of streptomycin (Invitrogen). HIV Rev-dependent GFP indicator Rev-A3R5-GFP cells were cultured in RPMI-1640 plus 10% FBS supplemented with 1 mg/ml geneticin (G418) (Sigma-Aldrich), 1 µg/ml puromycin (Sigma-Aldrich), and 100 units/ml of penicillin and 100 µg/ml of streptomycin (Invitrogen).

### Plasmid transfection and virus production

HIV-1 DNA vectors, pHIV-1(NL4-3) and pHIV-1(89.6), were obtained from the NIH AIDS Reagent Program. The env-defective pNL4-3 derivative pNL4-3/KFS and the HIV-1 Env expression vector, pNLΔΨEnv, were described previously ^51,53^. pCMV3-CD34 and the pCMV3-Empty vector were purchased from Sino Biological. The murine leukemia virus (MLV) packaging plasmid, pCL-Eco, was obtained from Addgene. The GFP-expressing retroviral vector pRetroQ-AcGFP1-N1 was purchased from Clontech. For HIV-1 virus production, HEK293T cells were co-transfected in 6-well plates with 1 µg of pHIV-1(NL4-3) plus the indicated doses of pCMV3-CD34 or pCMV3-Empty vector. Supernatants were harvested at 48 hours post co-transfection. HIV viruses were also assembled in 10-cm dishes by transfection of HEK293T cells with 10 µg of pHIV-1(NL4-3) or pHIV(89.6). Supernatant was collected at 48 hours post transfection. For MLV-GFP virus production, HEK293T cells were co-transfected in 6-well plates with 1 µg of pCL-Eco and 1 µg of pRetroQ-AcGFP1-N1 plus the indicated doses of pCMV3-CD34 or the pCMV3-Empty vector. Supernatants were collected at 48 hours post co-transfection. For HIV(KFS)(Env) virus production, HEK293T cells were transfected in 10-cm dishes with 5 µg of pNL4-3/KFS and 5 µg of pNLΔΨEnv, and the supernatant was harvested at 48 hours post transfection. For HIV-1 p24 release assays, cells were co-transfected with 1 μg of HIV-1(NL4-3) DNA and the indicated amounts of pCMV3-CD34 or pCMV3-Empty vector using Lipofectamine 2000 (Invitrogen). Supernatant was collected at 48 hours post transfection. To assemble the WNV pseudoviruses, HEK293T cells were co-transfected in 10-cm dishes with specific plasmid combinations, 10 μg of pCMV-WNV(NSW2011)-CprME (Virongy), and 10 μg of pWNVII-REP-G/Z (kindly provided by Dr. Theodore Pierson, NIAID). For VEEV pseudovirus assembly, HEK293T cells were co-transfected with 5 μg of pCMV-VEEV-Env, 5 μg of pCMV-VEEV-Cap, and 10 μg of pAlphaPRO-GFP (Virongy).

### Virus infectivity assays

For quantifying HIV-1 infectivity, HIV-1 particles were produced in the presence of pCMV3-CD34 or pCMV3-Empty vector. Particles were harvested at 48 hours, normalized by p24 content, and used to infect Rev-A3R5-GFP cells (0.2 to 0.5 million cells per infection). The percentage of GFP^+^ cells was quantified by flow cytometry at 48 to 72 hours post infection. For the MLV-GFP virus infectivity assay, NIH/3T3 cells were pretreated with Infectin (kindly provided by Virongy) for 1 hour as suggested by the manufacturer. Cells were infected with MLV-GFP virus in the presence of Infectin for 4 hours. Infected cells were washed and cultured in complete medium, and the percentage of GFP^+^ cells was quantified by flow cytometry at 72 hours post infection. To determine the inhibition of HIV-1 replication in CD34^+^ human CD4 T cells, Rev-A3R5-GFP cells (1 million cells) were electroporated with 500 ng of pCMV3-CD34 or pCMV3-Empty using a Cell Line Nucleofector Kit R (Lonza). After 2 days post electroporation, Rev-A3R5-GFP cells were infected with p24-normalized HIV-1(NL4-3) or HIV(KFS)(Env) virus. The percentage of GFP^+^ cells was quantified by flow cytometry at 48 to 72 hours post infection. HIV-1(NL4-3) and HIV-1(89.6) were used to infect Kasumi-3 cells. Briefly, cells were pretreated for 1 hour with Infectin (Virongy), and then infected in the presence of Infectin. After 4 hours, infected cells were washed and cultured in complete medium. Infection supernatants were harvested at days 0, 3, 5, 7, and 10 post infection for p24 ELISA.

To determine the infectivity of HIV-1 virus produced from CD34^+^ cord blood HSPCs, cells were cultured in StemSpan SFEM II (Stemcell) with or without expansion media (ExpM, with recombinant human cytokine mixture of SCF, Flt3L, and TPO at 100 ng/ml) (Stemcell). Cord blood HSPCs were also cultured in ExpM with or without recombinant human GM-CSF (R&D Systems) (100 ng/ml) plus recombinant human TNF-α (Biolegend) (2.5 ng/ml) for 3 to 7 days. Cells (1.5 million) were then infected with HIV-1(VSV-G) pseudotyped virus overnight, and then washed, and the media were replaced with fresh media. At 4 days post infection, virus supernatant was collected, and viral infectivity was quantified by spinoculation (300 x g, 2 h) of Rev-A3R5-GFP indicator cells with p24-normalized virus ^65^. The percentage of GFP^+^ cells was quantified at 8 days post infection by flow cytometry (FACSCalibur, BD Biosciences).

To measure HIV particle release from Kasumi-3 cells, 4 µg of pHIV-1(NL4-3) or pHIV(89.6) or pCMV3-Empty was electroporated into 2 million Kasumi-3 cells using Cell Line Nucleofector Kit R (Lonza). Electroporated cells were cultured at 37°C, 5% CO_2_, and virus supernatants were harvested at day 4, 7, and 9 post electroporation for p24 ELISA. Cells were also collected for surface staining.

To quantify the infectivity of HIV-1 virus particles produced from Kasumi-3 cells, cells were cultured with or without recombinant human GM-CSF (R&D Systems) (100 ng/ml) plus recombinant human TNF-α (Biolegend) (2.5 ng/ml) for 7 days. Cells (2 million) were then electroporated with HIV-1 DNA (4 µg) or HIV-1 DNA plus a CD34 expression vector (0.5 and 1 µg) using a Cell Line Nucleofector Kit R (Lonza). Following electroporation, cells were cultured either in medium or with GM-CSF plus TNF-α for 4 days. Virus supernatants were harvested, and virus infectivity was measured by infecting Rev-A3R5-GFP indicator cells with p24 normalized viruses. The percentage of GFP^+^ cells was quantified by flow cytometry (FACSCalibur, BD Biosciences).

To quantify the infectivity of HIV-1 virus produced from Kasumi-3 and CD34^ko^ Kasumi-3 cells, one million cells were infected with HIV-1 virus overnight, and cells were washed and the media were replaced with fresh media. At 4 days post infection, virus supernatants were collected, and viral infectivity was quantified by spinoculation of Rev-A3R5-GFP indicator cells (300 x g, 2h) with p24-normalized viruses. The percentage of GFP^+^ cells was quantified at 10 days post infection by flow cytometry (FACSCalibur, BD Biosciences). To evaluate the HIV-1 replication in Kasumi-3 and CD34^ko^ Kasumi-3 cells, HIV-1(NLENG1-ES-IRES) virus was also used to infect 0.5 million Kasumi-3 and CD34^ko^ Kasumi-3 cells by spinoculation (300 x g, 2 h). Cells were washed and replaced with fresh media. At 2 days post infection, viral infection was quantified by measuring GFP expression using flow cytometry (FACSCalibur, BD Biosciences).

To quantify HIV replication in Kasumi-3 and CD34^KO^-Kasumi-3 cells, equal p24 levels of HIV-1(NL4-3) particles (3 or 30 ng of p24) were used to infect (spinoculation, 300 x g, 2 h) Kasumi-3 or CD34^KO^-Kasumi-3 cells (1 million). Cells were washed, and cultured in fresh medium for two weeks. Supernatants were collected at 2, 4, 6, 8 and 10 days, and analyzed by an in-house HIV p24 ELISA kit.

To quantify the infectivity of WNV and VEEV pseudoviruses assembled in the presence or absence of CD34, HEK293T cells cultured in 12-well plates were infected with equal levels of pseudoviruses normalized by genomic RNA copies, and viral infectivity was measured by GFP expression at 2 days post infection.

### CD34 virion incorporation assay

To determine CD34 virion incorporation, HIV-1 particles assembled in the presence of pCMV3-CD34 or pCMV3-Empty at the indicated dosage were harvested at 48 hours post transfection in HEK293T cells. Viruses were filtered and purified by ultra-speed centrifugation through a 6%–18% OptiPrep gradient. CD34 and viral p24 proteins in each fraction were analyzed by western blot using antibodies against CD34 (Clone 563, BD Pharmingen) or HIV-1 p24 (mouse anti-HIV-1 p24 monoclonal antibody, clone 183-H12-5C, NIH AIDS Reagent Program). For the magnetic beads pull-down assay, HIV-1 particles were assembled in 6-well plates by co-transfection of HEK293T cells with pHIV-1(NL4-3) DNA (1 µg) plus pCMV3-CD34 or pCMV3-Empty vector at the indicated dosage. Particles were harvested at 48 hours post transfection, normalized for HIV-1 p24, and subjected to immuno-magnetic capture as previously described ^41^. Briefly, magnetic Dynabeads Pan Mouse IgG (Invitrogen) were conjugated with anti-human CD34 Antibody (Clone 563, BD Pharmingen) or purified Mouse IgG1, κ Isotype Ctrl Antibody (clone MG1-45, Biolegend) for 30 min at room temperature. After conjugation, the antibody-conjugated beads were incubated with p24 normalized CD34-incorporated viral particles for 1 hour at 37°C. The complex was pulled down with a magnet and washed with cold PBS twice. Captured viral particles were eluted in 10% Triton x-100 PBS, diluted, and quantified by p24 ELISA. To determine whether CD34 incorporation affects the incorporation of the HIV envelope, HIV-1 particles produced in the presence of pCMV3-CD34 or pCMV3-Empty at the indicated dosage were harvested 48 hours post transfection in HEK293T cells. Particles were purified using a sucrose gradient-based low-speed virus concentration kit (Virongy). Particles were analyzed by western blot using antibodies against HIV gp41(human anti-HIV-1 gp41 antibody, clone 2F5, NIH AIDS Reagent Program, # ARP-1475) and HIV p24 (mouse anti-HIV-1 p24 monoclonal antibody, 183-H12-5C, NIH AIDS Reagent Program, # ARP-3537).

### Viral attachment assay

HIV virion particles produced in the presence of pCMV3-CD34 or pCMV3-Empty were incubated with Rev-A3R5-GFP cells (prechilled at 4°C for 1 hour) at 4°C for 2 hours. The cells were then washed extensively (5 times) with cold PBS buffer and then lysed with LDS lysis buffer (Invitrogen) for analysis by western blot.

Similarly, HIV or NL43/KFS virion particles produced in the presence of pCMV3-CD34 or pCMV3-Empty were incubated with Hela and Hela JC53 cells were prechilled at 4°C for 1 hour) at 4°C for 2 hours. The cells were then washed extensively (5 times) with cold PBS buffer and then lysed with LDS lysis buffer (Invitrogen) for analysis by western blot.

### Viral entry assay

HIV-1 entry assay was performed as previously described ^66^. Briefly, viruses were generated by co-transfection of HEK293T cells with three plasmids: pHIV-1(NL4-3), pAdvantage (Promega) and pCMV4-3BlaM-Vpr (kindly provided by Dr. Warner C. Greene) at a ratio of 6:1:2. Supernatant was harvested at 48 hours post transfection, concentrated, and then used for infection of Rev-A3R5-GFP cells that had been electroporated with pCMV3-CD34 or pCMV3-Empty. β-lactamase and CCF2 (LiveBLAzer™ FRET-B/G Loading Kit with CCF2-AM, Invitrogen) measurements were performed using a 407-nm violet laser with emission filters of 525/50 nm (green fluorescence) and 440/40 nm (blue fluorescence), respectively. Green and blue emission spectra were separated using a 505LP dichroic mirror. The UV laser was turned off during the analysis. Flow cytometry was performed using a Becton Dickinson LSR II (Becton Dickinson). Data analysis was performed using FlowJo software (FlowJo, BD).

### Western blots

Cells were lysed and denatured by boiling in LDS lysis buffer (Invitrogen), subjected to SDS-PAGE, transferred to nitrocellulose membrane, and then incubated overnight at 4°C with one of the following primary antibodies: Purified mouse anti-human CD34 (Clone 563, BD Pharmingen) (1:1000 dilution), mouse anti-HIV-1 p24 monoclonal antibody (183-H12-5C, NIH AIDS Reagent Program) (1:1000 dilution), human anti-HIV-1 gp41 antibody (2F5, NIH AIDS Reagent Program) (1:1000 dilution), or anti-GAPDH goat polyclonal antibody (Abcam) (1:1000 dilution). Membranes were incubated with an anti-mouse IgG, HRP-linked antibody (Cell Signaling) (1:2000 dilution) for 60 min at room temperature or with HRP-labeled anti-goat IgG (H+L) antibody (KPL) (1:2500 dilution) for 30 min at room temperature. Chemiluminescence signal was detected by using West Femto chemiluminescence substrate (Thermo Fisher Scientific), and images were captured with a CCD camera (FluorChem 9900 Imaging Systems) (Alpha Innotech, San Leandro, CA, USA).

To confirm the CD34 knockout at the protein level, Kasumi-3 cells were pelleted, washed with 1x DPBS, and lysed with 1x LDS sample buffer (Thermo Scientific). The samples were sonicated with a microtip sonicator (Qsonica) (20A, 5s on/off) for 1 min, boiled, analyzed with Bis-Tris Mini Protein Gels (4–12%) (ThermoScientific), and then transferred onto nitrocellulose membrane (ThermoScientific). The blots were further blocked with 5% skim milk (Difco) in 1x TBST, washed, and incubated with an mouse anti-human CD34 antibody (BD) (1:1000), and then an HRP-conjugated horse anti-mouse IgG (1:1000) (Cell Signaling). The blots were also probed with goat anti-GAPDH polyclonal antibodies (Abcam) (1:2000) followed by an HRP-conjugated anti-goat IgG (H+L) antibody (KPL) (1:2000 dilution). The blots were imaged using chemiluminescent substrate, ECL (Thermo Scientific), and images were acquired using a Bio-Rad ChemiDoc imaging system and analyzed with the Bio-Rad Image Lab software.

### HIV p24 ELISA

Detection of extracellular HIV-1 p24 was performed using an in-house p24 ELISA kit. Briefly, microtiter plates (Sigma-Aldrich) were coated with anti-HIV-1 p24 monoclonal antibody (183-H12-5C, NIH AIDS Reagent Program). Samples were incubated for 2 hours at 37°C, followed by washing and incubating with biotinylated anti-HIV immune globulin (HIVIG, NIH AIDS Reagent Program) for 1 hour at 37°C. Plates were washed and incubated with avidin-peroxidase conjugate (R&D Systems) for 1 hour at 37°C. Plates were washed and incubated with tetramethylbenzidine (TMB) substrate and kinetically read using an ELx808 automatic microplate reader (Bio-Tek Instruments, Winooski, VT, USA) at 630 nm.

### Surface staining

For CD34 surface staining, 0.5 to 1 million HEK293T cells or Rev-A3R5-GFP cells were stained with mouse anti-human CD34 (Clone 563, BD Pharmingen) followed by staining with PE-cyanine5-labeled F(ab’)2-goat anti-mouse IgG (H+L) secondary antibody (Invitrogen). For CD4, CXCR4, CD34 surface staining on Kasumi-3 cells, 0.5 million cells were stained with mouse anti-human CD34 (Clone 563, BD Pharmingen), mouse anti-human CD184 (Clone 12G5, BD Pharmingen), or mouse anti-human CD4 (Clone RPA-T4, BD Pharmingen) followed by staining with AF488-labeled goat anti-mouse antibody (Invitrogen). Kasumi-3 cells were also stained with mouse anti-human CD162 (BD Pharmingen, clone KPL-1), mouse anti-human CD43 (BD Pharmingen, clone 1G10), mouse anti-human CD34, (BD Pharmingen, clone 563), human podocalyxin antibody (R&D Systems, clone 222328), human endoglycan/PODXL2 antibody (R&D Systems, clone 211805), TMEM123 monoclonal antibody (Invitrogen, clone 297617), anti-human CD164 antibody (Biolegend, clone 67D2), human TIM-1/KIM-1/HAVCR antibody (R&D Systems, clone#219211), mouse anti-human MUC1 (CD227) (BD Pharmingen, Clone HMPV), or human MUC-4 antibody, (R&D Systems, Clone 781631). Following primary antibody staining, cells were also stained with Alexa Fluor 488 goat anti-mouse IgG secondary antibody (2 mg/ml, Invitrogen) (1:10 dilution). Kasumi-3 cells were also stained with FITC mouse anti-human CD195 (BD Pharmingen, #555992) and FITC mouse IgG2α, k isotype (BD Pharmingen, #555573) to measure CCR5.

To determine the effect of GM-CSF/TNF-α on the expression of SHREK proteins on Kasumi-3 cells, 1 million Kasumi-3 cells were stimulated either with 0.1% BSA or with GM-CSF/TNF-α (100 ng/ml of GM-CSF plus 2.5 ng/ml of TNF-α) at every alternate day for 10 days. Kasumi-3 cells were stained with mouse anti-human CD34 (clone#563) (BD Biosciences), mouse anti-human CD162 (PSGL-1) (clone#KPL-1) (BD Biosciences), anti-human CD164 (clone#67D2) (Biolegend), mouse anti-human CD43 (clone#1G10) (BD Biosciences), or a mouse IgG1, κ Isotype antibody (clone#MG1-45) (Biolegend), followed by staining with an Alexa Fluor 488 goat anti-mouse IgG secondary antibody (2 mg/ml) (Invitrogen).

To identify whether HIV viral proteins can downregulate CD34 expression, HEK293T cells were co-transfected in a 6-well plate with 100 ng pCMV3-CD34 plus the indicated doses of pHIV-1(NL4-3), pPA-Gag, an HIV-1 Vpu expression vector (NIH AID Reagent Program, Cat#10076), an HIV-1 Nef expression vector (NIH AID Reagent Program, Cat#6454), or pCMV3-Empty vector using Lipofectamine 3000 (Thermo Fisher). After 48 h post transfection, cells were stained with a mouse anti-human CD34 antibody (Clone#563) (BD Pharmingen), followed by staining with an Alexa Fluor 488-labeled goat anti-mouse secondary antibody (Invitrogen).

Primary human bone marrow CD34^+^ cells and cord blood human stem and progenitor cells (HSPCs) were stained with a FITC-labelled mouse anti-human CD34 monoclonal antibody (clone 4H11[APG])(abcam) or a FITC-labelled mouse IgG1 antibody (clone B11/6) (abcam). Cord blood human stem and progenitor cells were also stained with a PE/Cy5-labelled anti-human CD4 antibody (clone RPA-T4) (Biolegend), a PE/Cy5-labelled mouse IgG1 antibody (clone MOPC-21) (Biolegend), a PE/Cy5-labelled anti-human CXCR4 antibody (clone 12G5) (Biolegend), or a PE/Cy5-labelled mouse IgG2a antibody (clone MOPC-173) (Biolegend, 400218).

### Generation of CD34 knockout cells using CRISPR-Cas9

The Kasumi-3 cells (1 million) were centrifuged and washed with 1x DPBS and electroporated with the ribonuclear complex (RNP) containing 100 µM CD34 sgRNA (Stemcell) and 2 µg Trucut Cas9 protein (Thermo Scientific) using the nucleofection kit R (Lonza) and Nucleofector II system (Amaxa) per the manufacturer’s recommendation. After 24 hours, electroporated cells were centrifuged and re-suspended into fresh complete RPMI-1640 medium supplemented with penicillin/streptomycin (100 U/ml) (GIBCO). The cells were grown for two weeks with frequent addition of fresh complete 1x RPMI medium every 4-5 days. Knockout cells were purified by magnet bead-based antibody removal of residual CD34^+^ cells. Briefly, CD34^-^Kasumi-3 cells were purified by negative depletion using an anti-human CD34 antibody (BD) and Dynabeads Pan Mouse IgG as previously described ^67^. The collected CD34^-^ cells were grown in RPMI-1640 medium and purified again to attain high purity of CD34^-^ cells.

## Supplemental figures

**Figure S1.**
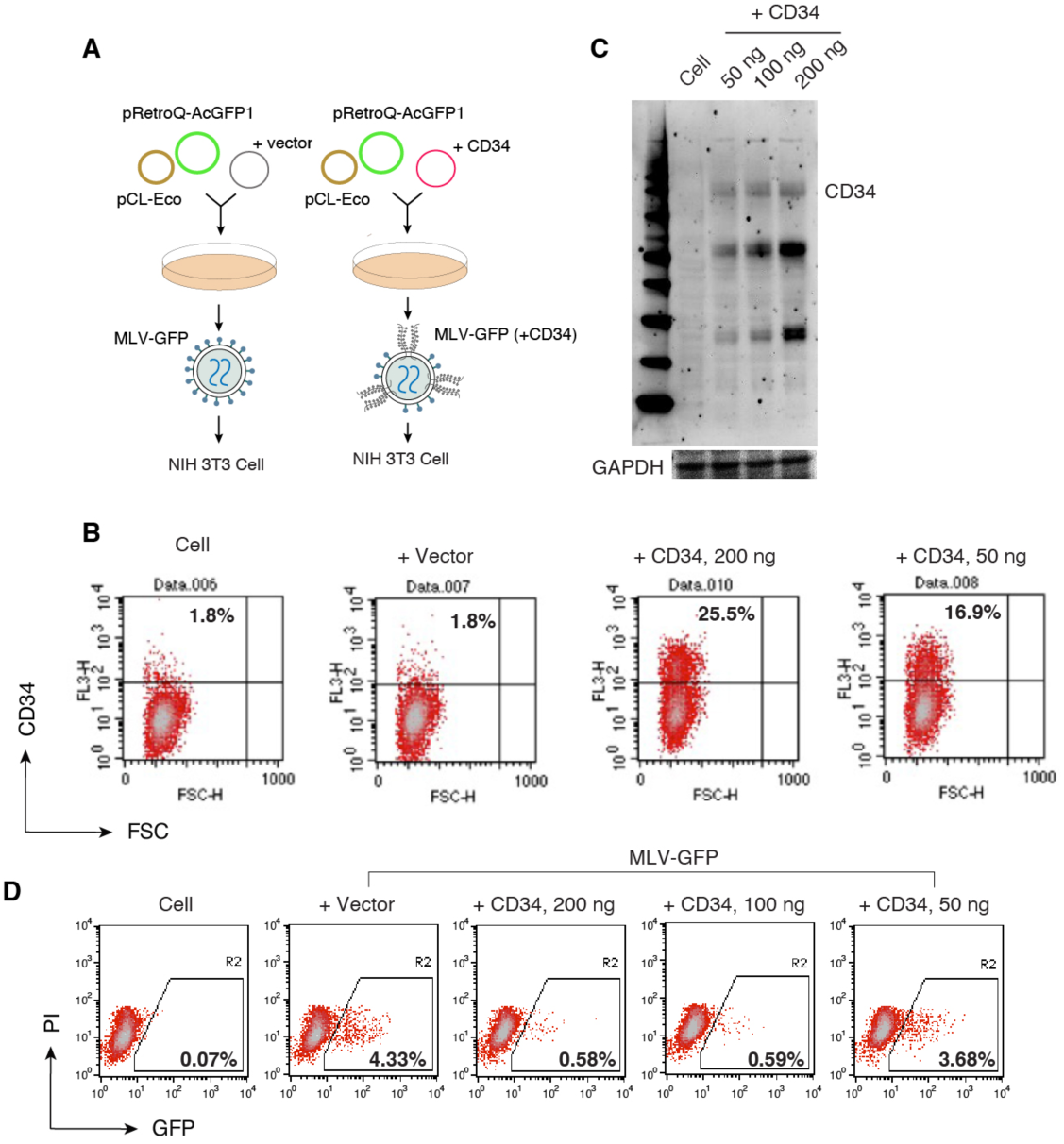
CD34 restricts MLV virus infectivity. (**A**) Schematic of the assay performed to determine effects of CD34 on MLV infectivity. (**B** and **C**) HEK293T cells were co-transfected with pCL-Eco and pRetroQ-AcGFP-N1 plus a CD34 expression vector or an empty control vector. Following co-transfection, expression of CD34 in cells was validated by surface staining of CD34 and flow cytometry (**B**) or by western blot (**C**). (**D**) MLV virions were harvested, and viral infectivity was quantified by infecting NIH 3T3 cells and measuring GFP expression.

**Figure S2.**
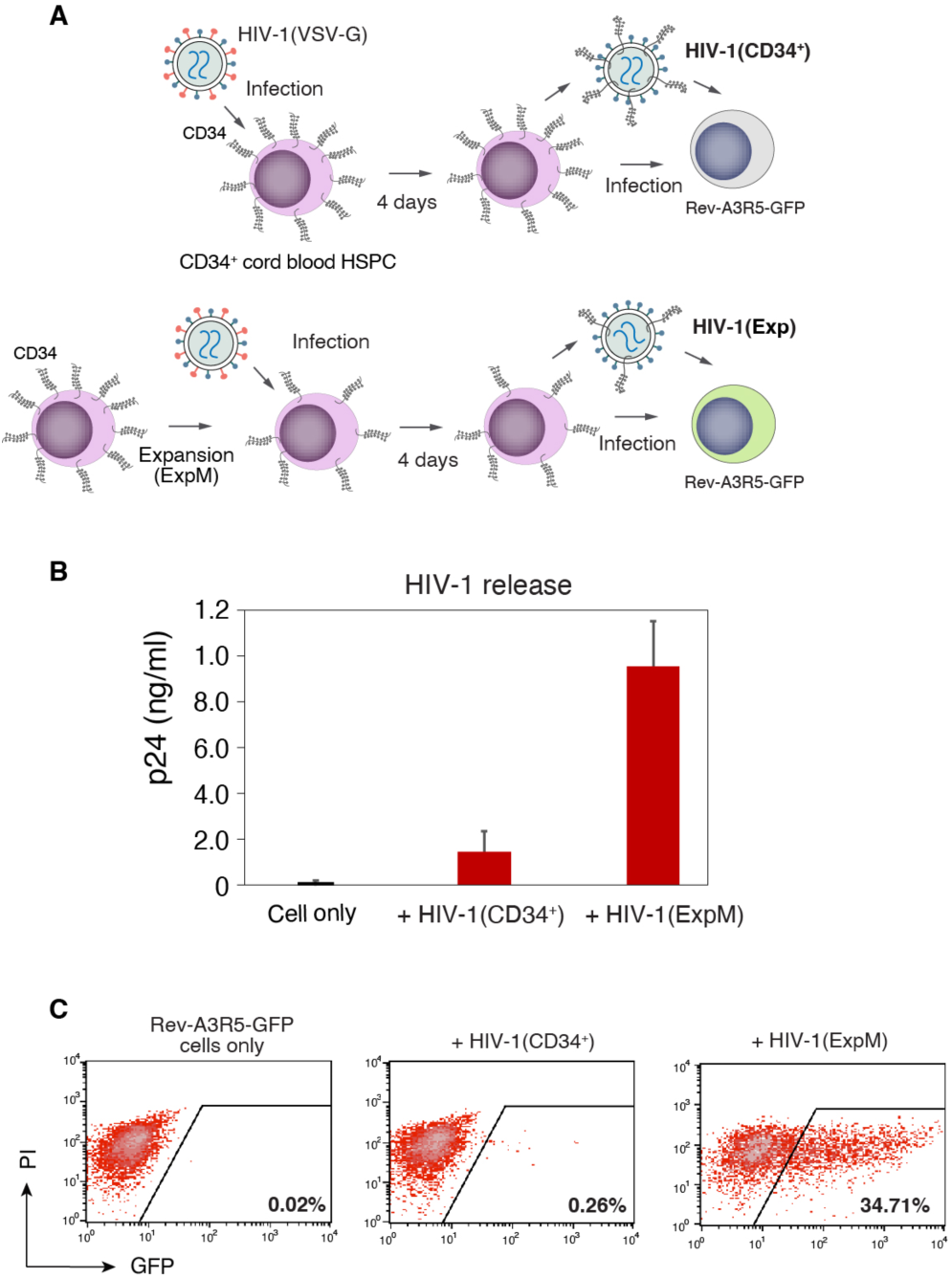
Direct infection of CD34^+^ cord blood HSPCs led to low-level release of non-infectious HIV particles. **(A)** Schematic of the infection procedures to determine HIV particle release and virion infectivity. (**B**) CD34^+^ cord blood HSPCs were either directly infected with HIV-1(VSV-G) or, for a comparison, cultured in expansion medium (ExpM) for 3 days, and then infected with HIV-1(VSV-G). Culturing CD34^+^ cord blood HSPCs in ExpM led to partial downregulation of CD34 (Fig. 4F). Following HIV-1(VSV-G) infection for 4 days, virion release was quantified with HIV p24 ELISA. (**C**) Quantification of virion infectivity. Viral particles released were harvested at 4 days post infection, and virus infectivity was quantified by infecting Rev-A3R5-GFP indicator cells, using an equal p24 viral input. GFP expression was quantified at 8 days post infection.

**Figure S3.**
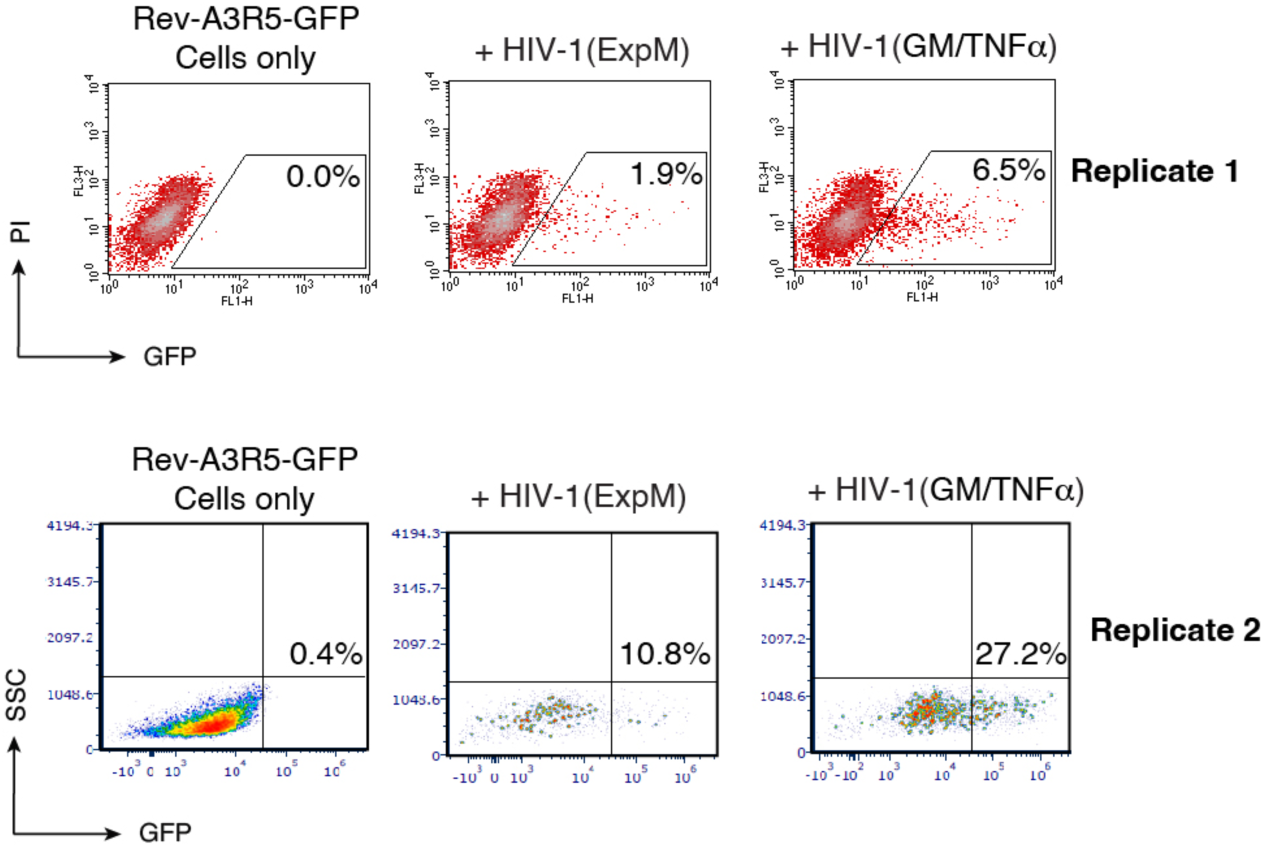
GM-CSF/TNF-α-mediated downregulation of CD34 enhances the infectivity of HIV progeny virions released from CD34^+^ cord blood hematopoietic stem/progenitor cells (HSPCs). Cells were cultured in expansion medium (ExpM) or in ExP plus GM-CSF/TNF-α for 7 days, and then infected with HIV-1(VSV-G) virus. At 4 days post infection, viruses released were harvested and virion infectivity was quantified by infecting Rev-A3R5-GFP indicator cells using an equal level p24 viral input. The percentage of GFP^+^ cells was quantified on 8 days post infection by flow cytometry. Shown are results from two independent experimental replicates.

**Figure S4.**
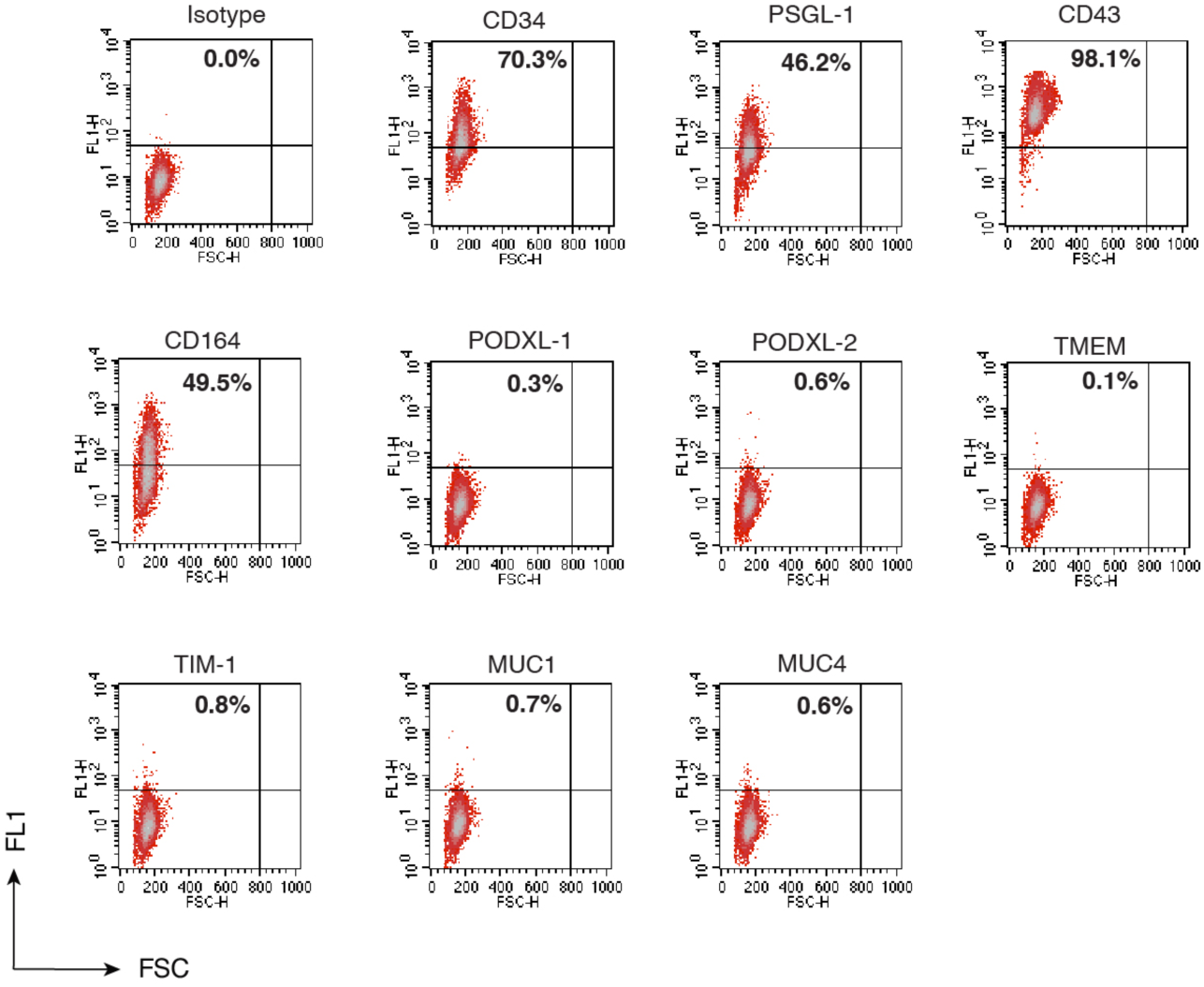
Expression of CD34 and other mucin-like SHREK proteins on Kasumi-3 cells. Kasumi-3 cells were stained with antibodies against CD34, PSGL-1, CD43, CD164, PODX-1, PODXL-2, TMEM, TIM-1, MUC1, or MUC4 and then analyzed by flow cytometry. Kasumi-3 cells expressed CD34, PSGL-1, CD43, and CD164, but not PODXL-1, PODXL-2, TMEM, TIM-1, MUC1, or MUC4.

**Figure S5.**
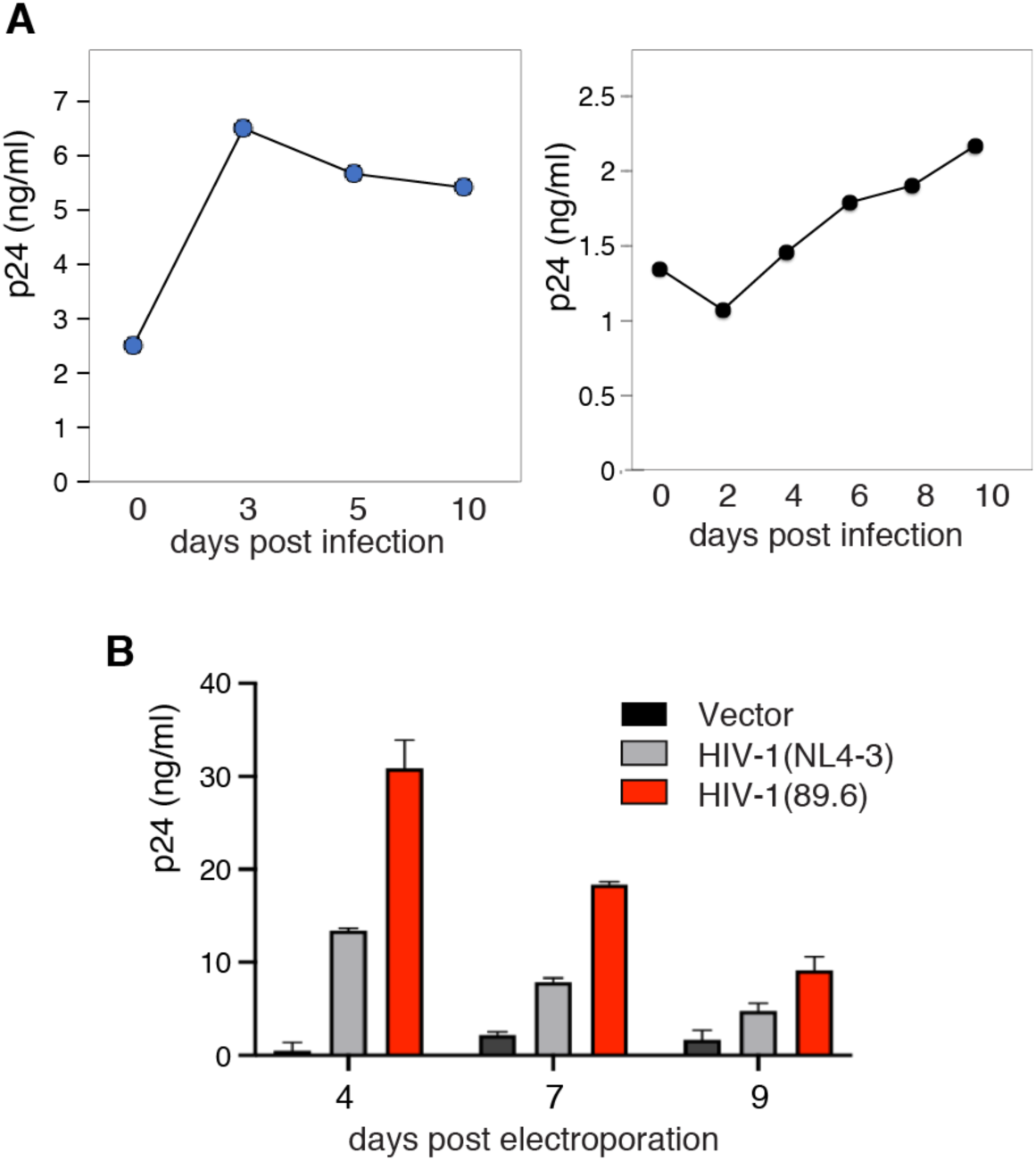
HIV-1 infection of Kasumi-3 cells. **(A)** Cells were infected with HIV-1(89.6) for 3.5 h, washed, and then cultured for 10 days. Viral p24 release was quantified by ELISA using cell culture supernatant. (**B**) Kasumi-3 cells were electroporated with HIV-1(NL4-3) DNA, HIV-1(89.6) DNA, or a control empty vector (4 μg). Viral replication was quantified by p24 release using ELISA of the cell culture supernatant.

**Figure S6.**
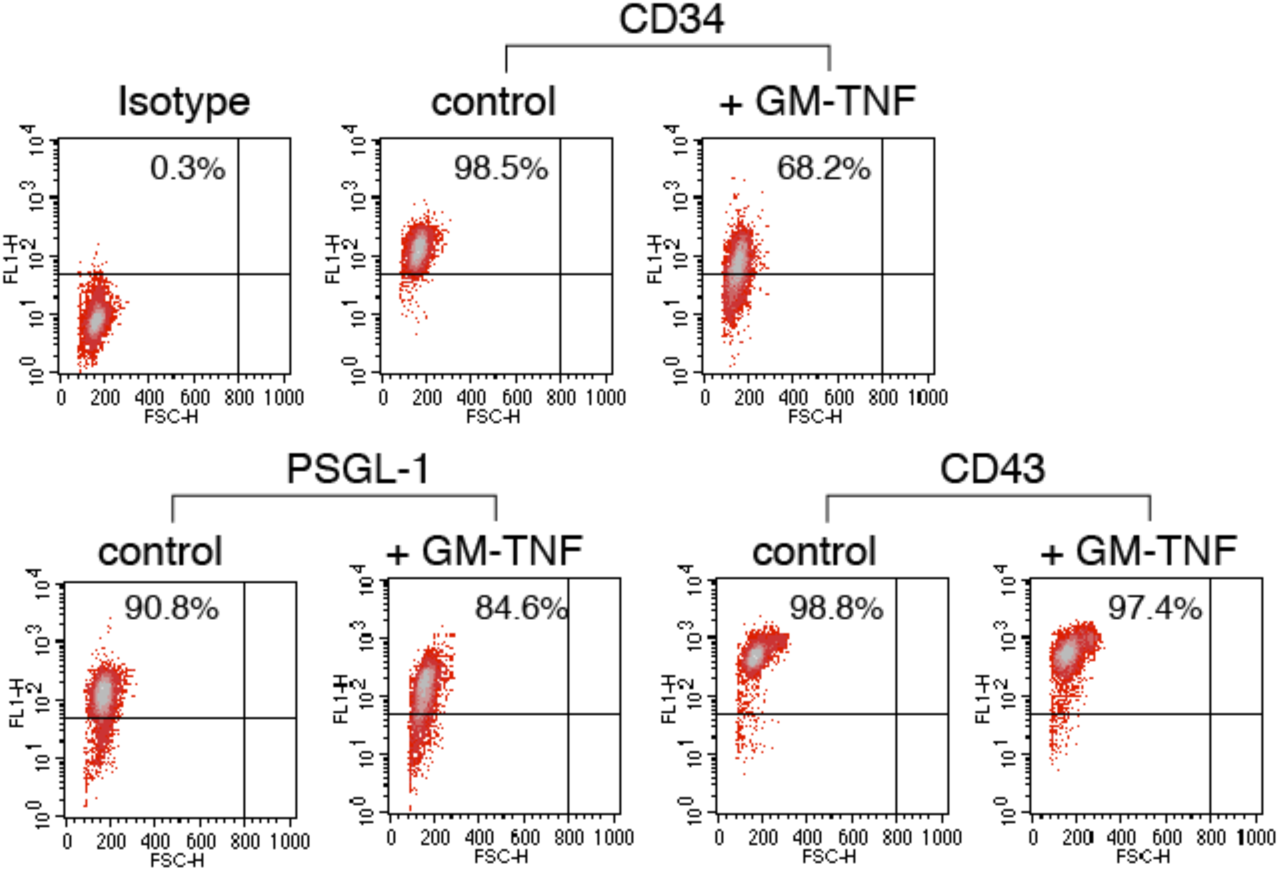
GM-CSF/TNF-α-mediated downregulation of CD34 but not PSGL-1 and CD43. Kasumi-3 cells were cultured in GM-CSF plus TNF-α or complete medium for 10 days, and surface expression of CD34, PSGL-1, and CD43 was measured by surface staining with antibodies against CD34, PSGL-1, or CD43 and then analyzed by flow cytometry. Shown are density plots of CD34, PSGL-1, and CD43 expression on Kasumi-3 cells cultured with or without GM-CSF plus TNF-α.

**Figure S7.**
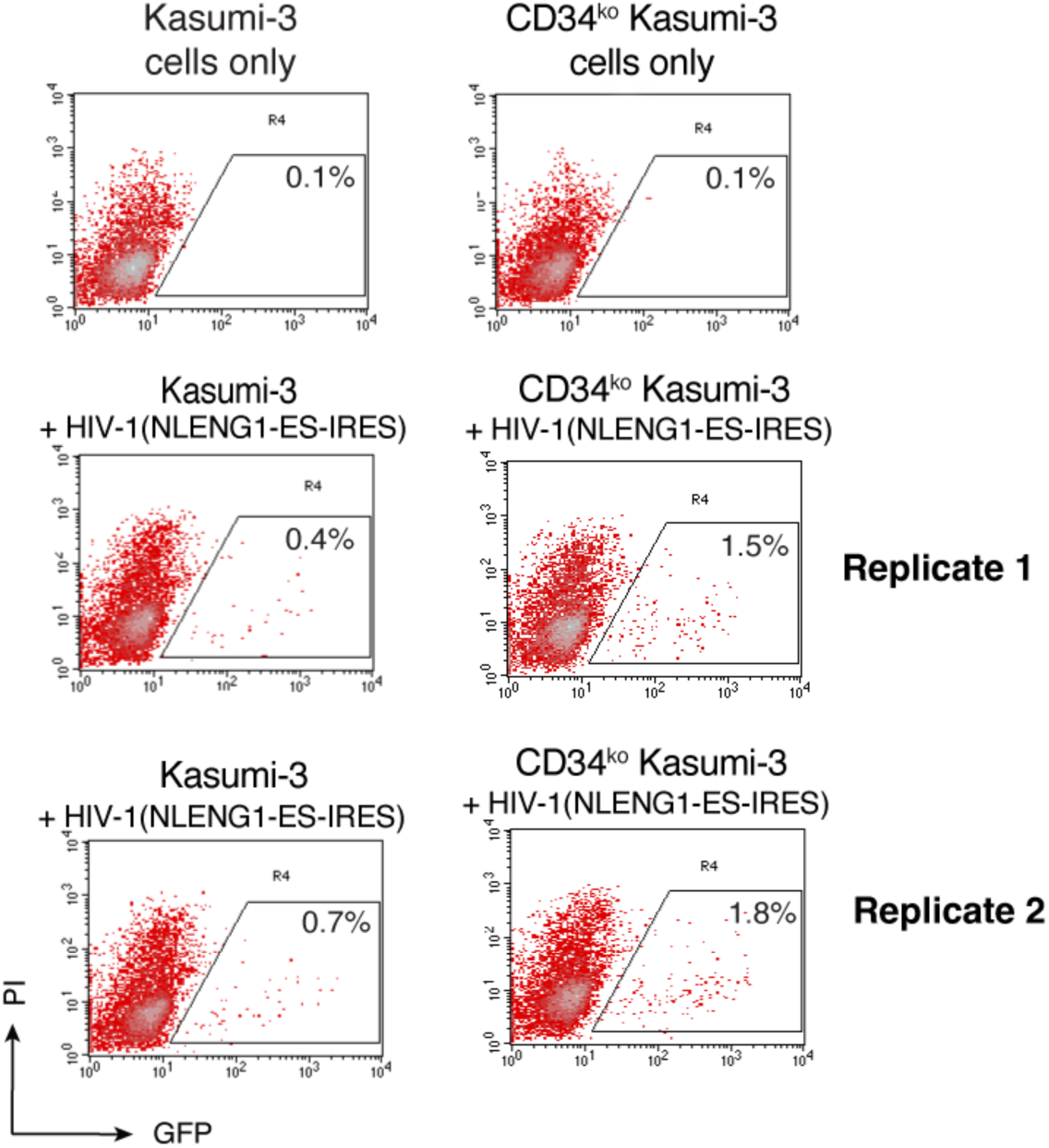
CRISPR/Cas9 knockout of CD34 leads to enhanced HIV replication in CD34^ko^. Kasumi-3 cells. Kasumi-3 and CD34^ko^ Kasumi-3 cells were identically infected with HIV-1(NLENG1-ES-IRES) using an equal p24 viral input. At 2 days post infection, viral replication was quantified by measuring GFP expression. Shown are results from two independent experimental replicates.

**Figure S8.**
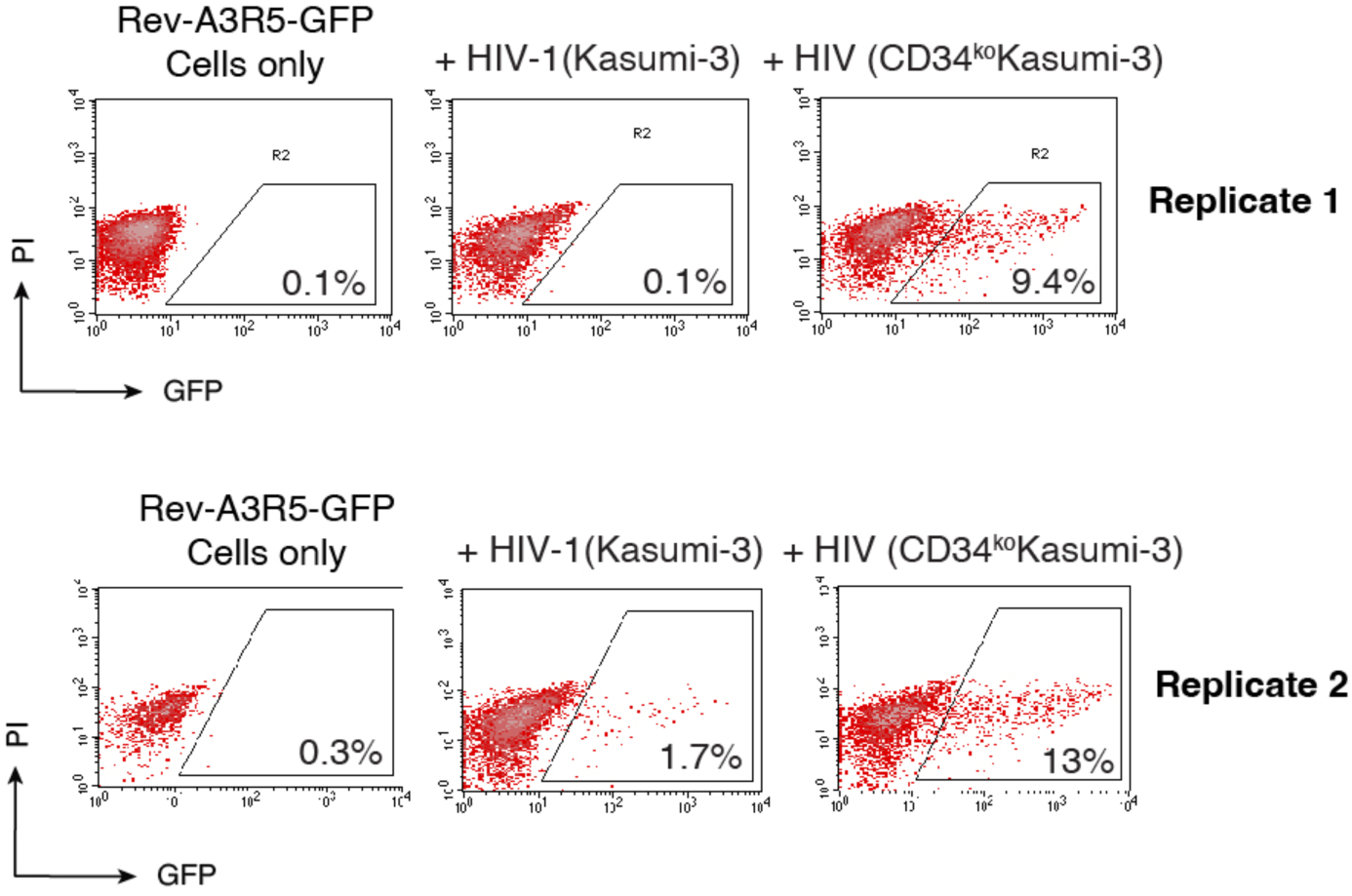
Enhanced infectivity of HIV-1 virions released from CD34^ko^ Kasumi-3 cells. Kasumi-3 or CD34^ko^ Kasumi-3 cells were infected with HIV-1 virus. At 4 days post infection, viruses released were harvested and virion infectivity was quantified by infecting Rev-A3R5-GFP indicator cells using an equal p24 viral input. The percentage of GFP^+^ cells was quantified at 10 days post infection by flow cytometry. Shown are results from two independent experimental replicates.

**Figure S9.**
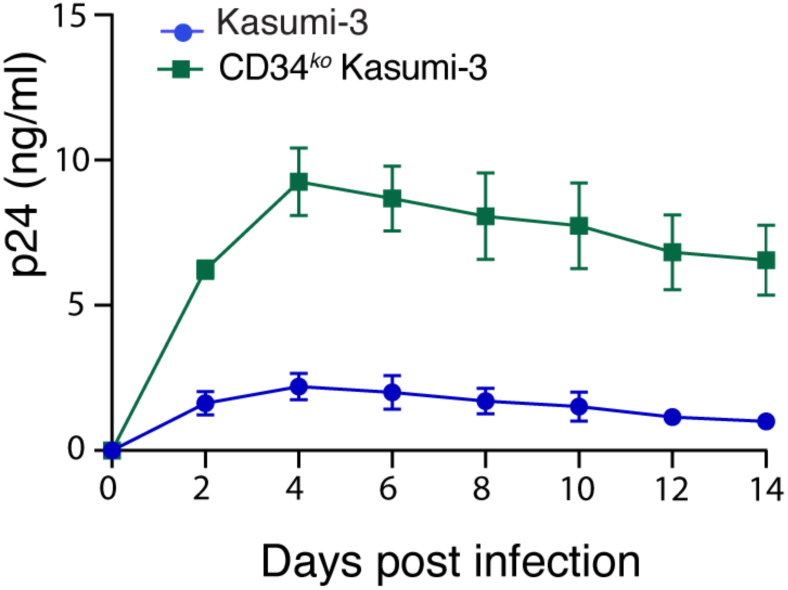
CRISPR/Cas9 knockout of CD34 leads to enhanced HIV replication in CD34^ko^. Kasumi-3 cells. Kasumi-3 and CD34^ko^ Kasumi-3 cells were identically infected with HIV-1(NL3-4) using an equal p24 viral input. HIV replication was quantified by measuring HIV p24 levels in the culture supernatant.

**Figure S10.**
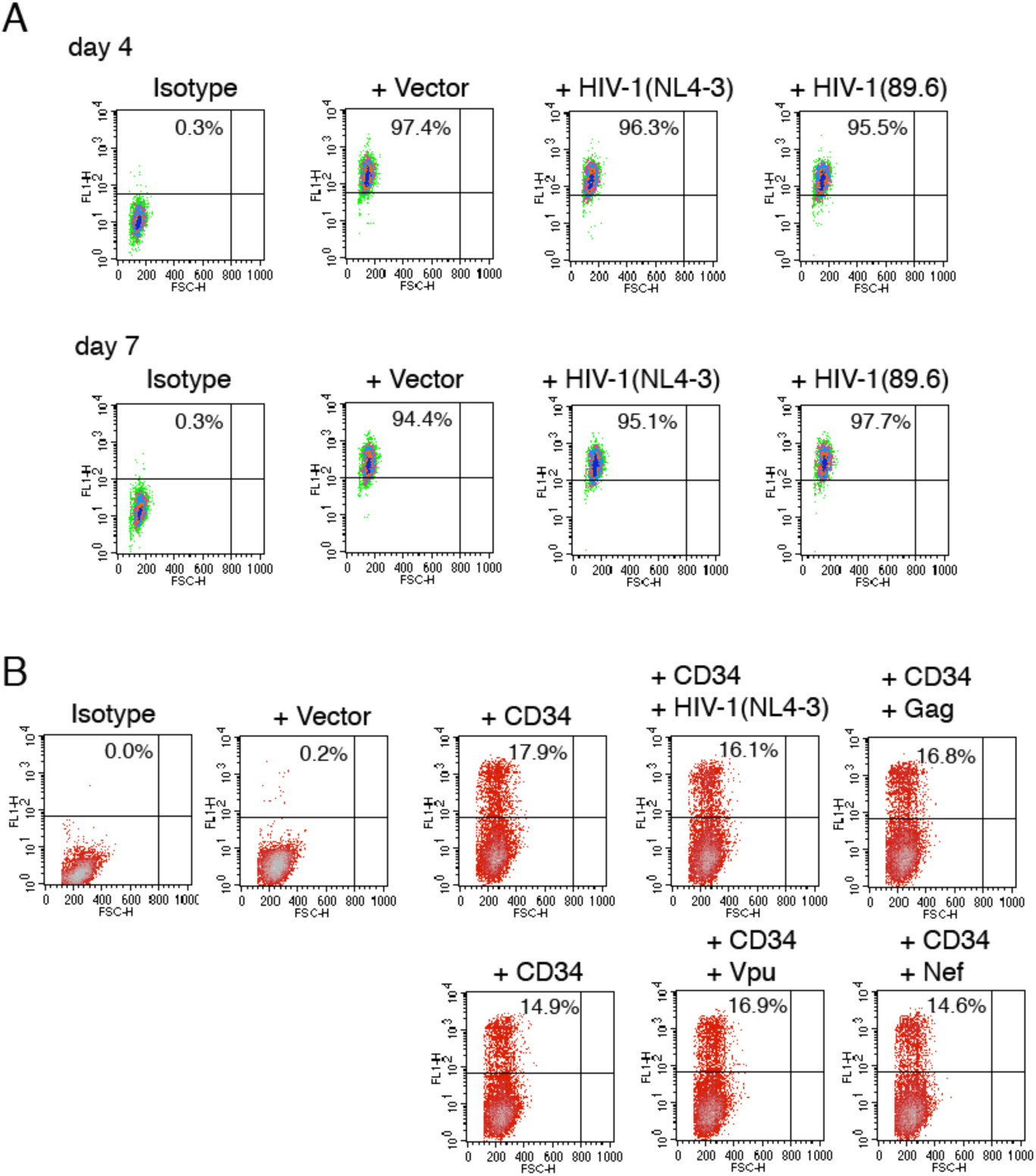
HIV-1 and viral Vpu, Nef, and Gag proteins do not downregulate CD34. **(A)** Kasumi-3 cells were electroporated with HIV-1(NL4-3) DNA, HIV-1(89.6) DNA, or a control empty vector (4 μg). Downregulation of surface CD34 was quantified at 4 and 7 days post electroporation. (**B**) HEK293T cells were transfected with a control empty vector or a CD34 expression vector (100 ng). Cells were also co-transfected with a CD34 expression vector plus HIV-1(NL4-3) DNA, a Gag expression vector, a Vpu expression vector, or a Nef expression vector (2 μg of each vector). Surface CD34 expression was quantified with flow cytometry at 48 h post co-transfection.

